# Label free autofluorescence imaging permits comprehensive and simultaneous assignment of cell type identity and reveals the existence of airway secretory cell associated antigen passages (SAPs)

**DOI:** 10.1101/2022.11.01.514675

**Authors:** Viral S Shah, Jue Hou, Vladimir Vinarsky, Jiajie Xu, Charles P Lin, Jayaraj Rajagopal

## Abstract

The specific functional properties of a tissue are distributed amongst its component cell types. The various cells act coherently, as an ensemble, in order to execute a properly orchestrated physiologic response. Thus, modern approaches to dissect physiologic mechanism would benefit from an ability to identify specific cell types in live tissues and image them in real time. Current techniques require the use of fluorescent genetic reporters that are not only cumbersome, but which only allow the simultaneous study of 2 or 3 cell types. We report a non-invasive imaging modality that capitalizes on the endogenous autofluorescence signatures of the metabolic cofactors NAD(P)H and FAD. By marrying morphological characteristics with autofluorescence signatures, all seven of the airway epithelial cell types can be distinguished simultaneously in real time. Furthermore, we find that this methodology for direct cell type specific identification avoid potential pitfalls with the use of ostensibly cell type-specific markers that can be altered by clinically relevant physiologic stimuli. Finally, we utilize this methodology to interrogate real-time physiology using a clinically relevant model of cholinergic stimulation and identify dynamic secretory cell associated antigen passages (SAPs) that are highly reminiscent of previously reported goblet cell associated antigen passages (GAPs) in the intestine.

**eLife’s Review Process:** eLife works to improve the process of peer review so that it more effectively conveys the assessment of expert reviewers to authors, readers and other interested parties. In the future we envision a system in which research is first published as a preprint and the outputs of peer review are the primary way research is assessed, rather than journal title.

Our editorial process produces two outputs: i) an assessment by peers designed to be posted alongside a preprint for the benefit of the readers; **i**) detailed feedback on the manuscript for the authors, including requests for revisions and suggestions for improvement.

Therefore we want to change how we construct and write peer reviews to make themuseful to both authors and readers in a way that better reflects the work you put into reading and thinking about a paper.

eLife reviews now have three parts:

- An **evaluation summary** (in two or three sentences) that captures the major conclusions of the review in a concise manner, accessible to a wide audience.
- A **public review** that details the strengths and weaknesses of the manuscript before you, and discusses whether the authors’ claims and conclusions are justified by their data.
- A set of private **recommendations for the authors** that outline how you think the science and its presentation could be strengthened.

All three sections will be used as the basis for an eLife publishing decision, which will, as always, be made after a consultation among the reviewers and editor. Each of the **public reviews** will be published (anonymously) alongside the preprint, together with a response from the authors if they choose. In the case of papers we reject after review, the authors can choose to delay posting until their paper has been published elsewhere.

If this is your first time going through this new process, we ask that you take some time to read our [Reviewer Guide](https://submit.elifesciences.org/cgi-bin/main.plex?form_type=display_rev_instructions%23writing), which discusses how we see each section will be used, what it should contain, and what we hope it accomplishes. And we remind you that, with the shift of reviews from private correspondence to public discourse, it is more important than ever that reviews are written in a **<u>clear and constructive manner</u>** appropriate for a public audience and mindful of the impact language choices might have on the authors.

## Introduction

The airway epithelium is comprised of myriad distinct cell types, many of which have only been described recently (Deprez et al., 2020; Goldfarbmuren et al., 2020; Montoro et al., 2018; Plasschaert et al., 2018). Studying the function of each of these cells and their contribution to the physiology of the entire epithelial ensemble often requires cell type-specific genetic perturbation. Indeed, even the simple identification of a given class of cells within a live tissue requires genetic labeling with a fluorophore. Thus, experimentally following the behavior of multiple different cell types simultaneously requires further prohibitive genetic modifications.

Currently, cells are usually identified post-fixation using immunohistochemical and immunofluorescence staining, but this prohibits assessing real time cellular dynamics and real time physiology. Since post-fixation cell identification is destructive, it also precludes the study of long-term processes or the effect of multiple serial stimuli which are both essential features of normal organ physiology. Additionally, markers used for cell type identification in unperturbed tissue can often be non-specific or misleading after injury (Lin et al., 2022) and has led to errors in lineage assignment. Finally, post-hoc identification of cell types also limits the study of rare cells or stochastically appearing cells, as the relevant cell types may not even be present in the selected experimental tissue or their markers are unstable or even unknown. We reasoned that a methodology for identifying and tracking diverse cell types within an organ in real time in a non-destructive fashion would permit the study of physiologically relevant processes that are currently hidden from view.

Label free imaging based on endogenous fluorescence of metabolic cofactors has been deployed to measure cell metabolism in various tissues, particularly in tumors and immune cells (Katz et al., 2002; Skala et al., 2007). NADH, NADPH, and FAD, the major endogenous fluorophores, have been used to monitor metabolism *in vitro* and *in vivo* (Gil et al., 2019; Heikal, 2010; Huang et al., 2002; Kasischke et al., 2004; Rocheleau et al., 2004; Sepehr et al., 2012; Tiede et al., 2007). Since NADH and NADPH cannot be distinguished by their autofluorescence, the combination has been reported as an aggregate quantity, NAD(P)H (Schaefer et al., 2019). To reduce the effects of light scattering, blood contamination and variation in number of mitochondria, a metabolic ratio of FAD/(NAD(P)H + FAD) is often employed (Chance et al., 1979). By exciting and detecting the fluorescence emission of these endogenous flurophores with 2-photon excited fluorescence microscopy (TPEF), the metabolic ratio can be measured in intact tissues and *in vivo* (Dilipkumar et al., 2019; Klinger et al., 2012; Kreiß et al., 2020). Indeed, in one study common intestinal epithelial cell types could be distinguished based on their overall morphology (Kreiß et al., 2020). Most recently these measures of metabolic activity have been used to identify immune cells in differing functional states during FACS *ex vivo* (Lemire et al., 2022). In this manuscript, we describe a live tissue explant system that permits label free cell type identification of both common and very rare epithelial cell types. Such precise cell type identification requires an assessment of the fine features of their autofluorescence signatures coupled to their morphology. We also use live autofluorescence imaging to reveal the existence of secretory cell associated antigen passages (SAPs) in the airway. The biology of such structures has only recently been dissected in the gut where they have been termed goblet cell associated antigen passages (GAPs) (Gustafsson et al., 2021; Gustafsson & Johansson, 2022; Knoop et al., 2014; McDole et al., 2012; Noah et al., 2019). Furthermore, we describe the real time dynamics of the formation of these structures for the first time using physiologic cholinergic stimulation of airway explants. Finally, we demonstrate that autofluorescence signature can be used to assign cell identity more specifically than some of the functional markers that are often currently used to describe disease pathology.

## Results

### Label free autofluorescence-based live imaging of an intact murine tracheal explant

We hypothesized that we would be able to specifically assign airway epithelial cell type identity by directly imaging specific metabolic signatures at single cell resolution. In order to preserve subtle and complex features of cellular and tissue architecture, we developed a tracheal explant setup that allows upright 2-photon imaging in a physiologic chamber over days (Figure 1A and Figure 1 Supplement 1A). To facilitate optimal detection and separation of NAD(P)H and FAD, the samples were scanned at 730nm and 900nm sequentially (Huang et al., 2002). A short pass 505nm dichroic was used to separate the NAD(P)H and FAD signal, and each fluorescence signal was detected with 450/100nm (NAD(P)H) and 540/80nm (FAD) band pass filters respectively (Supplement 1B). Using this setup, we ascertained NAD(P)H and FAD autofluorescence emissions across the excitation spectrum in the tracheal explant model (Figure 1B). Due to the curvature of the trachea, in order to visualize the epithelial surface in similar planes required digital “flattening” during image analysis where the sub-epithelial region was linearized to the same Z-plane. To accomplish this, we utilized second harmonic generation (SHG) imaging of collagen fibers to identify the basement membrane and developed a MATLAB algorithm to “flatten” the epithelial surface (Appendix 1). The SHG signal was excited with 900nm laser.

**Figure 1.**
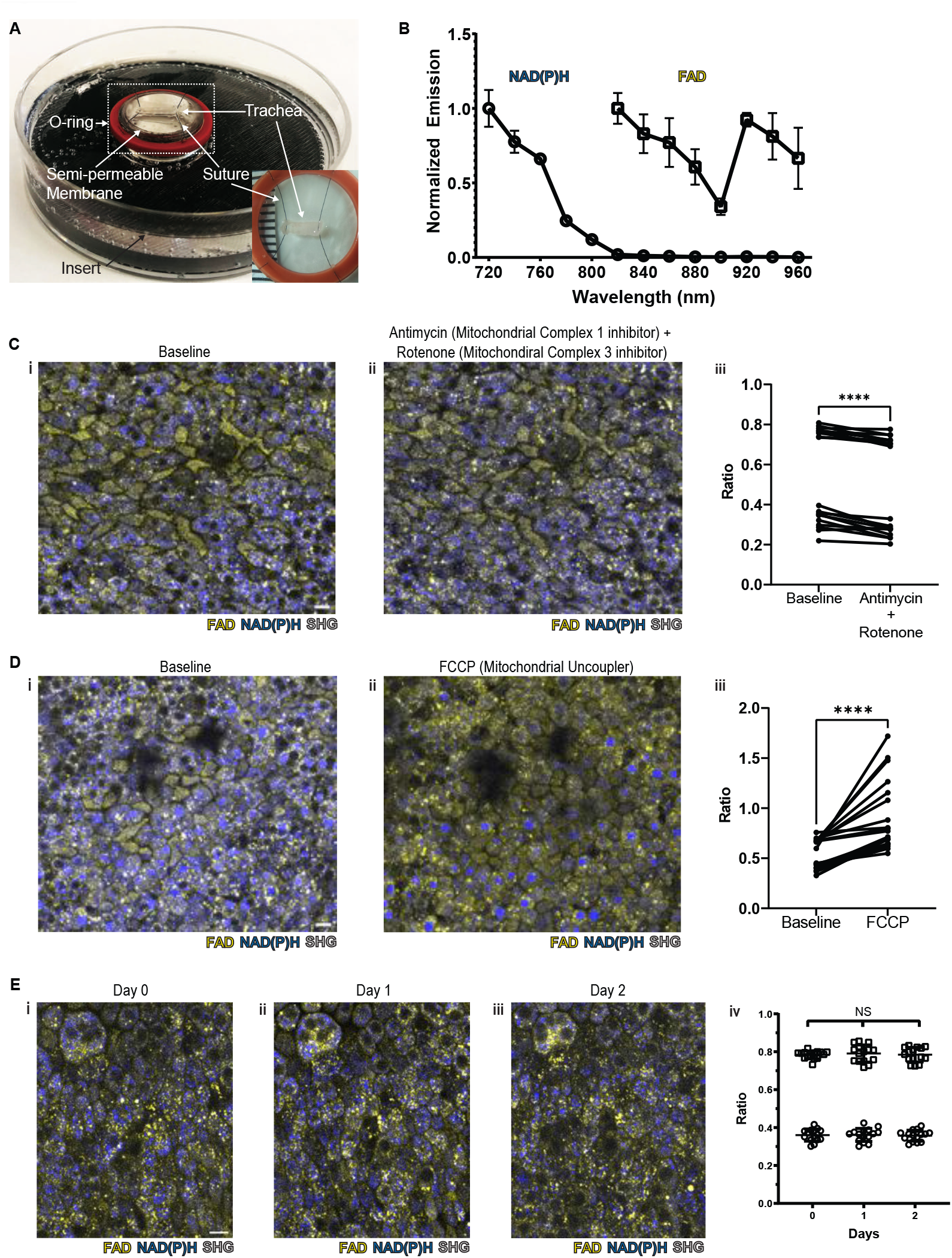
NAD(P)H and FAD autofluorescence can be measured from murine tracheal epithelial cells and is stable across days. (A) Mounting of trachea explant for upright 2-Photon imaging. Semipermeable membrane (6.5 mm Transwell®, 0.4 μm Pore Polyester Membrane Insert) was inverted and fixed to custom 3D printed insert in 60mm dish using silicone adhesive. Trachea was dissected and sutured onto o-ring for additional stability (Figure 1A inset and Figure 1 supplement 1A). Trachea and O-ring were placed onto the semipermeable membrane for upright imaging and explant culture. Inset showing a trachea sutured to O-ring in a zoomed view. (B) Autofluorescence emission normalized to maximal emission at different excitation wavelengths for both NADH and FAD. (C) NADH (blue), FAD (yellow), and second harmonic generation (SHG, gray) autofluorescence image of murine tracheal explant at (i) baseline, (ii) addition of antimycin (10uM - complex I inhibitor) and rotenone (1uM - complex III inhibitor) showing increased blue fluorescence, and (iii) quantification of autofluorescence ratio (FAD/FAD+NAD(P)H) with a decrease in ratio. Each point is a single cell. (D) baseline, (ii) addition of Carbonyl cyanide p-trifluoro-methoxyphenyl hydrazone (FCCP - mitochondrial uncoupler) showing decrease in blue fluorescence and increase in ratio (iii). Each point is a single cell. (E) Timecourse images of NAD(P)H (blue), FAD (yellow), and second harmonic generation (SHG, gray) of the same region on tracheal explant at day 0 (i), day 1 (ii), and day 2 (iii) showing similar fluorescence. (iv) Quantification of Ratio (FAD/FAD+NAD(P)H) of individual cells showing stability across 0, 1, and 2 days. ****P<0.0001 by paired t-test. Scale bars = 10 μm.

**Figure 1 Supplement 1.**
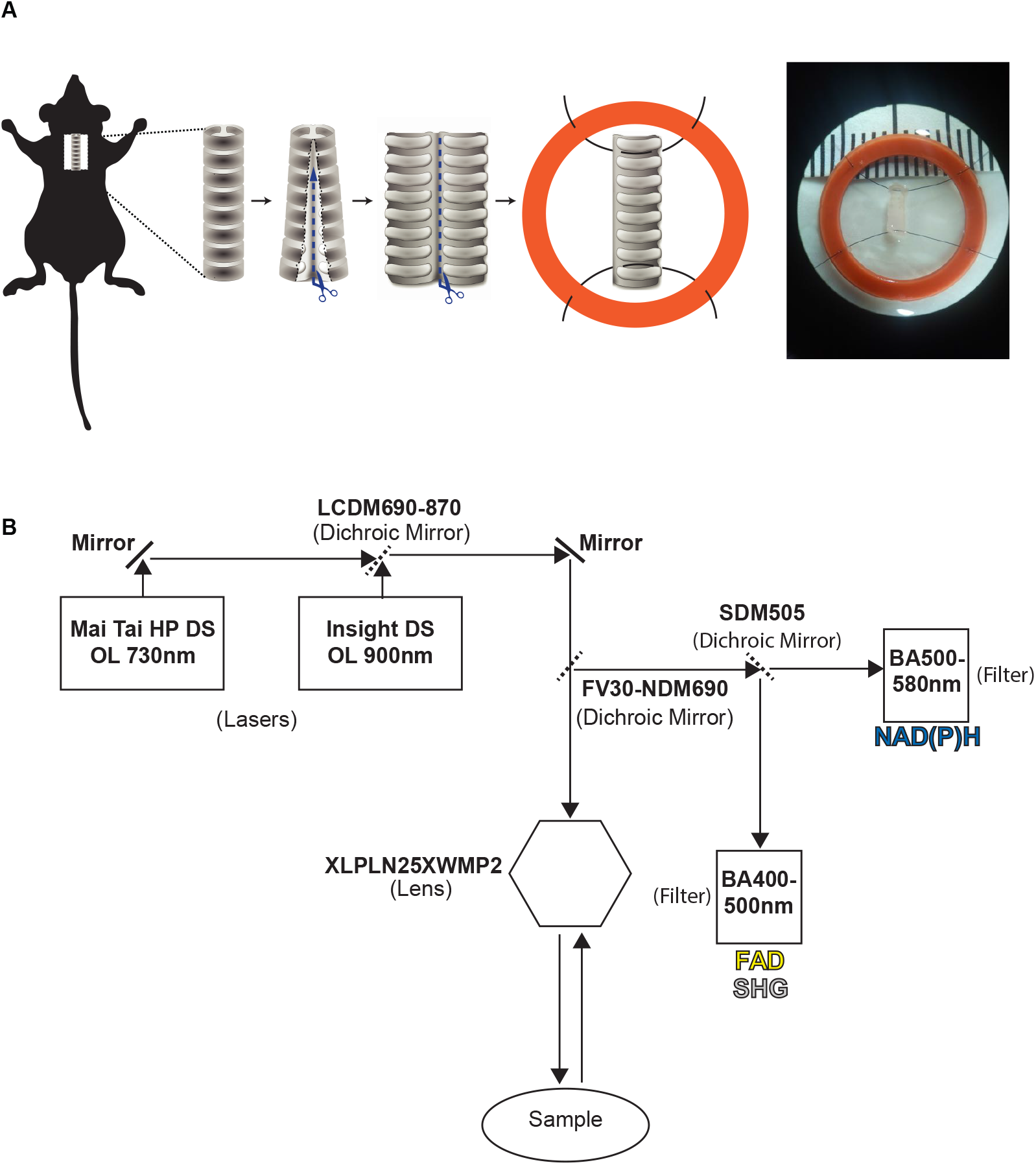
(A) Dissection and fixing trachea to O-ring with suture. Immediately after euthanasia, longitudinal excision was performed upon the tracheal tube after which further dissection along the posterior membrane separated the trachea into two pieces. One piece was consequently fixed to an O-ring using suture to prevent movement and allow securing of the trachea to the membrane insert. (B) Microscope filter and laser setup showing Insight laser set at 900nm and Mai-Tai at 730nm with a sharp cut dichroic mirror (SDM505) reflecting light less than 505 facilitated detection of FAD, second harmonic generation (SHG) and transmitting light greater that 505nm for and NAD(P)H detection.

To verify that the measured autofluorescence signals reflect cellular metabolism in real-time, we utilized pharmacologic agents to experimentally alter NAD(P)H and FAD. Treatment with rotenone and antimycin A, inhibitors of mitochondrial complex I and complex III, lead to a decrease in the FAD/(NAD(P)H + FAD) fluorescence ratio at the airway surface as predicted from similar experiments conducted in breast cancer cells (Figure 1C) (Hou et al., 2016).

Treatment with FCCP, a mitochondrial uncoupler, led to an anticipated increase in autofluorescence intensity ratio, as similarly previously documented (Figure 1D) (Hou et al., 2016). In order to determine signal stability, we measured the autofluorescence intensity ratio over 2 days of tracheal explant culture. The signal remained stable (Figure 1E).

### Autofluorescence imaging discriminates the three common airway epithelial cell types and demarcates airway epithelial structures and fine details of cell shape at the single cell level

As previous studies have been able to visualize common cell types in the intestinal epithelium using autofluorescence (Kreiß et al., 2020), we first optimized our label free imaging approach to identify the common airway epithelial cell types: ciliated cells, secretory cells, and basal cells. In order to register autofluorescence live images to that of fixed and stained images, we utilized characteristics of the second harmonic generation (SHG) from subepithelial collagen to identify fiduciary marks which facilitated exact localization of single cells. We first visualized ciliated cells with live autofluorescence imaging given their very unique morphology. As seen in Figure 2A, ciliated cells have a robust FAD signal which is primarily located near the cell surface in the location of cilia. Fixation and staining with acetylated tublin (ac-tub, green) demonstrated coincidence of this autofluorescence signature and the presence of ciliated cells identified by immunofluorescence. Secretory cells demonstrated a more robust NADH signal (Figure 2B) that was in sharp contrast to that of ciliated cells. Post fixation and staining with CCSP (magenta) demonstrates that autofluorescence intensity ratio can also define secretory cell borders. Similarly, basal cells can also be identified by their autofluorescence signatures (Figure 2C) and correspond to cells positive for keratin 5 (KRT5, cyan). These differences in autofluorescence ratio can be quantified for the individual cell types and can be used to distinguish ciliated, secretory, and basal cells (Figure 2D).

**Figure 2.**
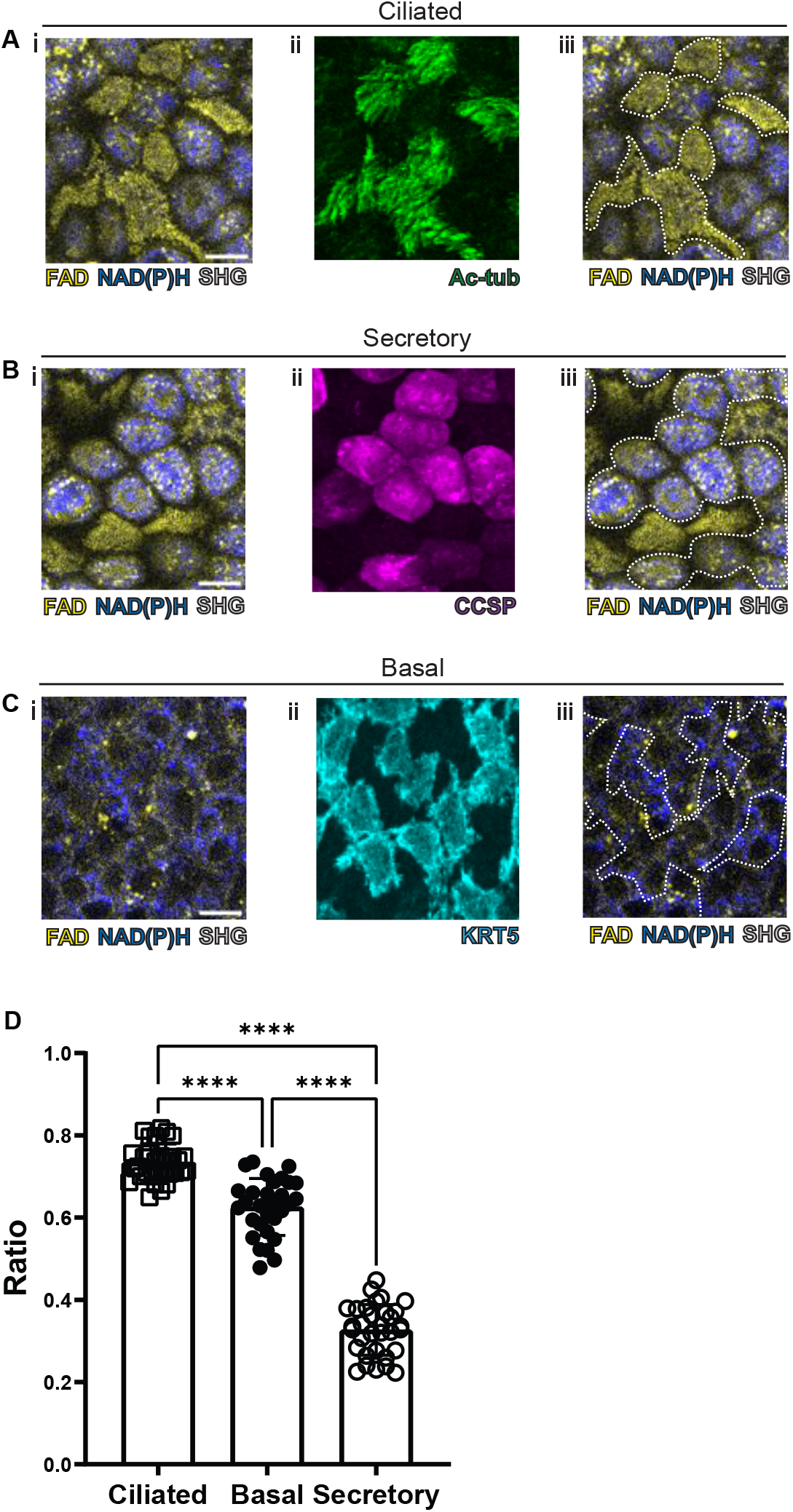
Autofluorescence imaging of common cell types. (A-C) NAD(P)H (blue), FAD (yellow), and second harmonic generation (SHG, gray) autofluorescence (i) and staining (ii) with acetylated tubulin (A) (ac-tub, green) (B) CCSP (magenta) and (C) KRT5 (cyan) for ciliated cells, secretory cells, and basal cells. Overlay of staining and autofluorescence (iii). (D) Quantification of FAD/FAD+NAD(P)H for ciliated cells, secretory cells, and basal cells, demonstrating distinct autofluorescence ratio. ANOVA test for multiple comparisons. **** p < 0.0001. Scale bars = 10 μm.

### Rare airway epithelial cell types can be distinguished using a combination autofluorescence signatures coupled to simple metrics of morphology

We next assessed the ability of our protocol to identify hillocks, a newly identified airway epithelial structure (Lin et al., 2022; Montoro et al., 2018) (Figure 3A). Due to the multilayered nature of hillocks, the structure is raised above the surrounding surface epithelia. The unique polygonal cell shape of hillock cells and discrete boundaries of hillocks are easily distinguishable with autofluorescence imaging. Post-fixation staining with keratin 13 (KRT13, magenta), a protein highly expressed by hillocks, demonstrates concordance with autofluorescence imaging.

**Figure 3.**
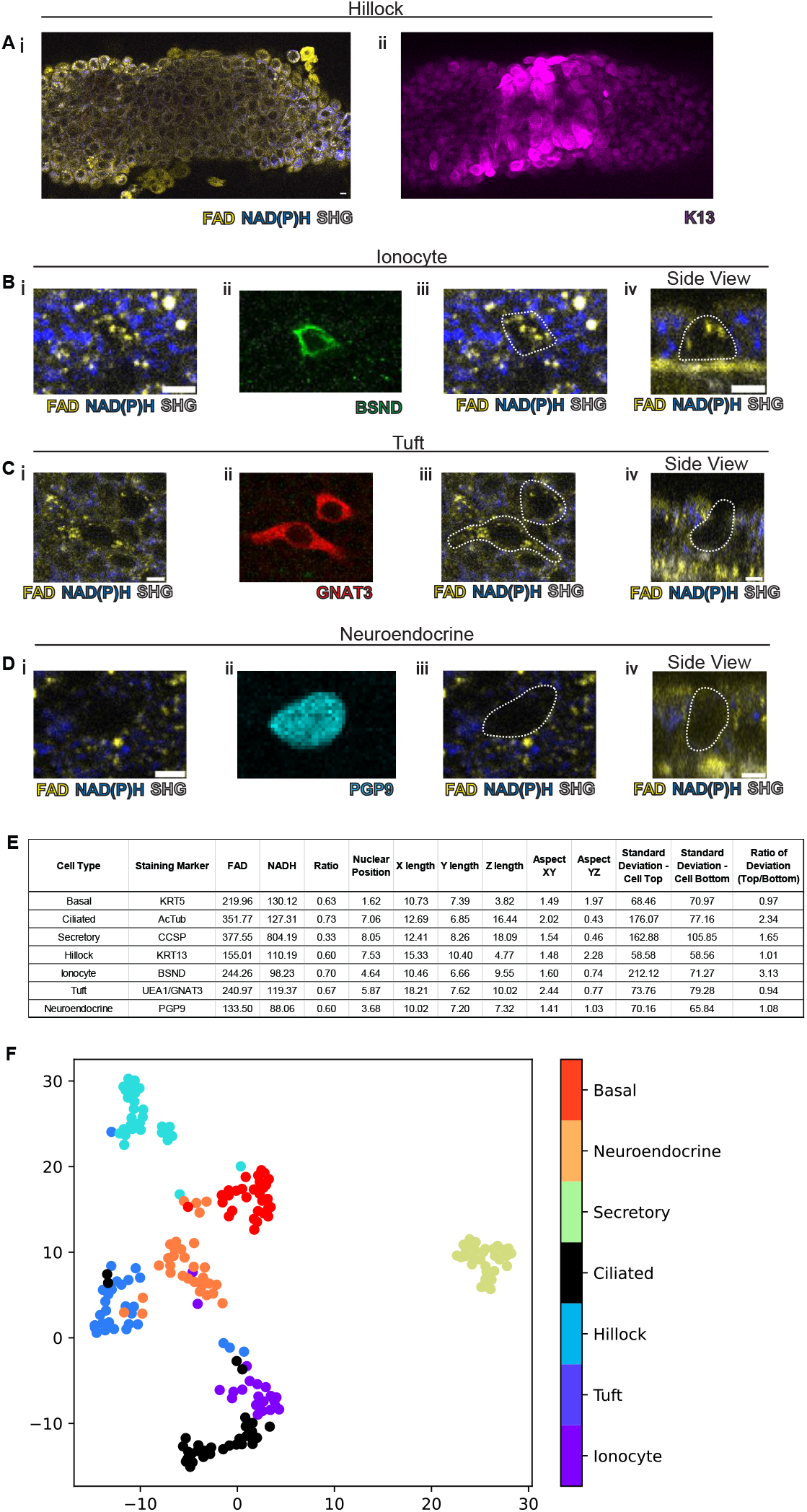
Rare cell detection using a combination of autofluorescence and unbiased clustering. NAD(P)H (blue), FAD (yellow), and second harmonic generation (SHG, gray) autofluorescence in (i) of (A) hillocks, (B) ionocytes, (C) tuft cells, (D) neuroendocrine cells. Staining in (ii) of hillocks (K13 - magenta), ionocytes (BSND - green), tuft cells (GNAT3 - red), neuroendocrine cells (PGP9 - cyan). Scale bars = 10 μm in (A) and 5 μm in (B-D). (iii) Overlay of cell outline from (ii) onto (i). Cross sectional imaging across the Z plane (iv) showing differential fluorescence in ionocytes. (E) Table with cell type-specific values of both autofluorescence (FAD, NADH, Ratio) and specific morphologic parameters. “X-length” = largest distance in X dimension. “Y-length” = distance in Y dimension. “Z-length” = largest distance in Z dimension. “Aspect XY” = Ratio of X-length / Y-length. “Aspect YZ” = Y-length / Z-length. “Standard deviation Cell Top” = standard deviation of autofluorescence ratio from above mid height. “Standard deviation Cell Bottom” = standard deviation of autofluorescence ratio from below mid height. “Ratio of standard deviation” is standard deviation cell top / standard deviation cell bottom. (F) Unbiased clustering from (E) demonstrating separate clusters for each cell type.

We then took on the much greater challenge of identifying the very rare cell types of the epithelium: pulmonary ionocytes, tuft cells, and neuroendocrine cells (Figure 3B-D). Ionocytes (Figure 3B) had a characteristic shape by autofluorescence which was similar to the shape of ionocytes delineated by immunohistochemical staining with BSND (green). Additionally, ionocytes were found to have puncta of high FAD signal towards the apical aspect of the cell (Figure 3Biv). The distinct shape of tuft cells could be identified by an autofluorescence ratio corresponding to immunofluorescent staining with GNAT3 (red). Of all airway epithelial cells, neuroendocrine cells displayed the lowest NAD(P)H and FAD autofluorescence and their location was confirmed with staining by PGP9 (cyan) (Figure 3D). Interestingly, as an aggregate, rare cells are characterized by very similar autofluorescence intensity ratios and could not be distinguished by this criterion alone (Figure 3E). Given the diversity of morphologies of airway cell types, we then quantified a number of morphometric measures to better distinguish rare cell identities (Figure 3 Supplement 1). We measured the distance of the nucleus from the basement membrane (nuclear position), the largest dimension in the X, Y, and Z plane (X length, Y length, Z length), the aspect ratios (X length / Y length) and (Y length / Z length), and the standard deviation of fluorescence ratio intensities across the cell in the Z plane (standard deviation of autofluorescence intensity ratio above the nucleus, standard deviation of autofluorescence intensity ratio below nucleus, and the ratio). We measured these parameters for both the common cells (basal, ciliated, and secretory), hillocks, and rare cells (ionocyte, neuroendocrine, and tuft). Aggregated data from a total of 30 basal cells, 32 ciliated cells, 32 secretory cells, 30 hillock cells, 21 ionocytes, 30 tuft cells, and 31 neuroendocrine cells were compiled (Figure 3E) with cell identities confirmed by immunofluorescence staining.

Given the spread of various morphologic variables across different cell types, we performed unsupervised clustering of the cell types based upon their autofluorescence intensity ratio and their morphological differences. In aggregate, these characteristics allowed the separation of all airway epithelial cell types into distinct clusters (Figure 3F, Figure 3 Supplement 2). We then trained three machine learning models using k-nearest neighbors, multinomial logistic regression, and xgboost algorithms to classify the different cell types. We utilized 75% of the total cells (154 cells) for machine learning training, leaving 25% (52 cells) for subsequent validation. We then compared the identity assigned by the machine learning algorithms based on the autofluorescence and morphology data to the known identity determined by staining markers. K-nearest neighbor analysis and multinomial logistic regression resulted in a 96% accuracy, while xgboost performed with 92% accuracy. This was suggestive of high-quality modeling with a Matthews Correlation Coefficient of 0.95, 0.95, and 0.91 (Figure 3 Supplement 3).

**Figure 3 Supplement 1.**
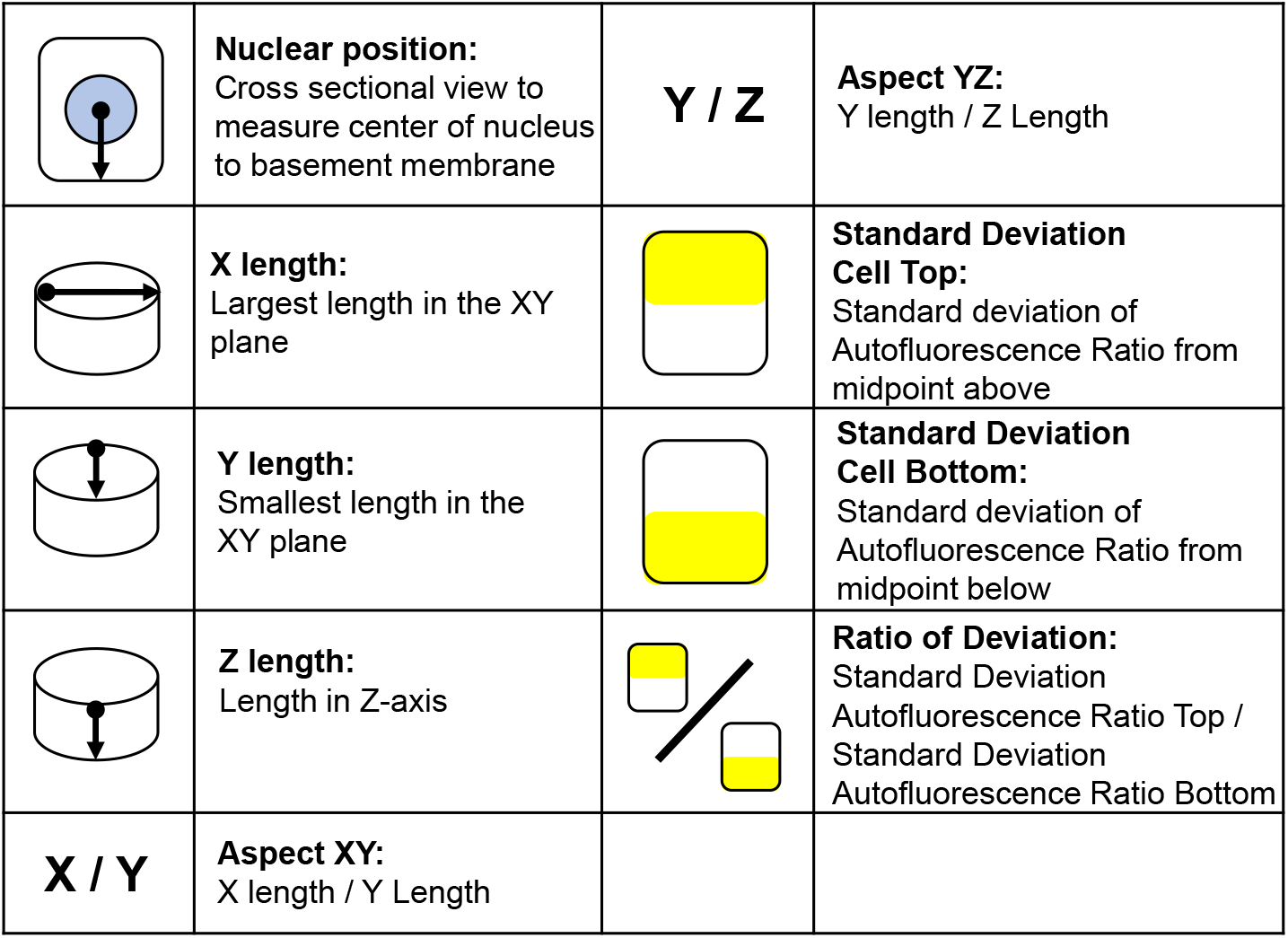
Description of morphologic analysis for all airway epithelial cell types.

**Figure 3 Supplement 2.**
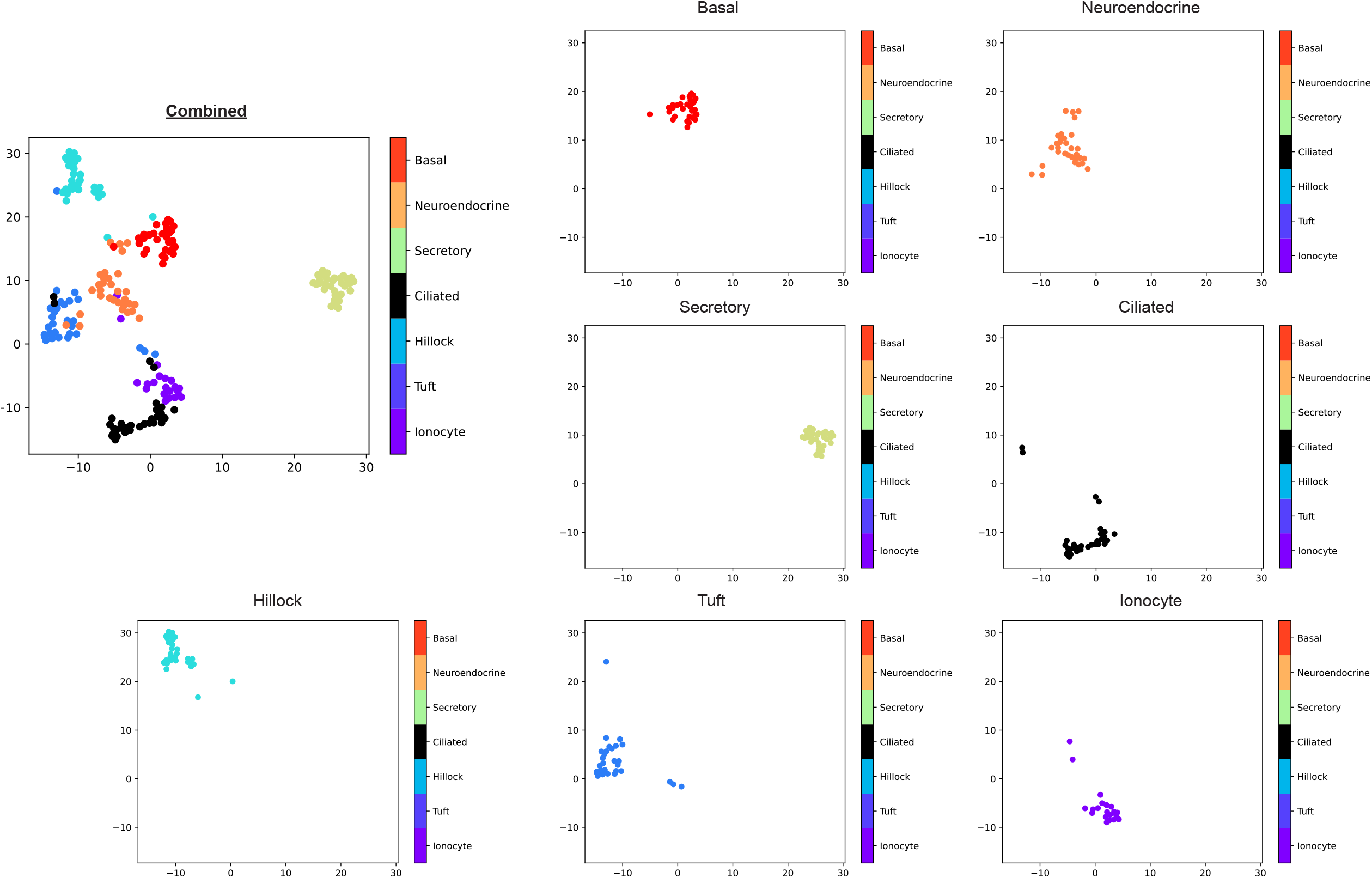
Individual groups of results of unsupervised clusters analysis of all airway epithelial cell types. Basal cells (red), Neuroendocrine cells (orange), Secretory cells (green), Ciliated cells (black), Hillock (cyan), Tuft (blue), Ionocyte (magenta).

**Figure 3 Supplement 3.**
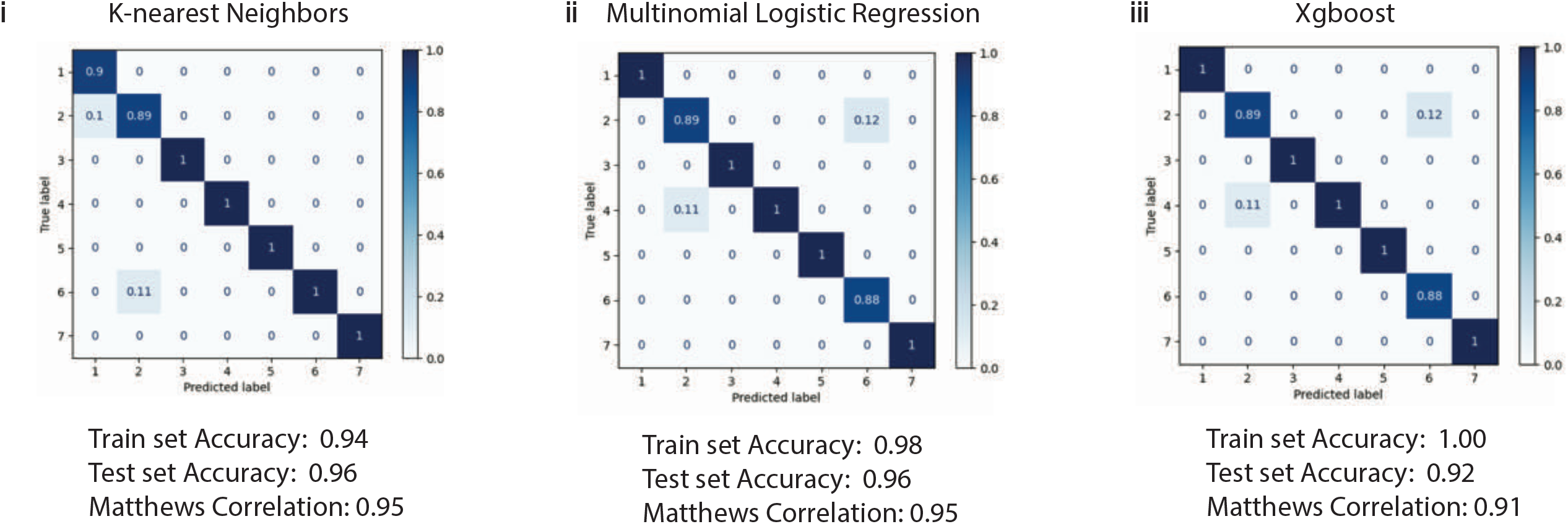
Confusion matrix demonstrating the performance of 3 algorithms on testing the clustering based on unsupervised clustering from Figure 3E-F. 152 cells were used for training and 52 cells were used to test each algorithm (i) k-nearest neighbors, (ii) multinomial logistic regression, and (iii) xgboost. The accuracy for training set of 152 cells was 0.94, 0.98, and 1.00 respectively. The accuracy for the test set of 52 cells was 0.96, 0.96, and 0.92 respectively. Matthews Correlation Coefficient is 0.95, 0.95, and 0.91 respectively. 1 = Ionocytes, 2 = Tuft cells, 3 = Hillock cells, 4 = Ciliated cells, 5 = Secretory cells, 6 = Neuroendocrine cells, 7 = Basal cells.

### Autofluorescence imaging can be used to interrogate real time physiology and reveals the existence of secretory cell associated antigen passages

In order to assess whether this technology is useful to assess real time physiologic parameters, we sought to employ a paradigmatic physiologic stimulus and assess whether the tissue response could be monitored dynamically. Cholinergic and adrenergic stimulation have historically been known to cause airway secretory cell granule release (Massaro et al., 1979; Yoneda, 1977). The club cell secretory protein, CCSP (also known as CC10, Uteroglobin, CC16, *SCGB1A1*) is a major constituent of these granules and is commonly used to assign secretory cell identity (Wong et al., 2009). We applied a classic cholinergic agonist, methacholine (which is used clinically to define airway hyper-responsiveness in asthma) to the airway explant model (Adler et al., 2013; Fischer et al., 2019; Webber & Widdicombe, 1987). Methacholine stimulation led to a 333% decrease in the CCSP immunofluorescence staining as measured by absolute fluorescence intensity (Figure 4A-B). Indeed, this stimulus-induced change in CCSP leads to a dramatically reduced ability to identify secretory cells.

**Figure 4.**
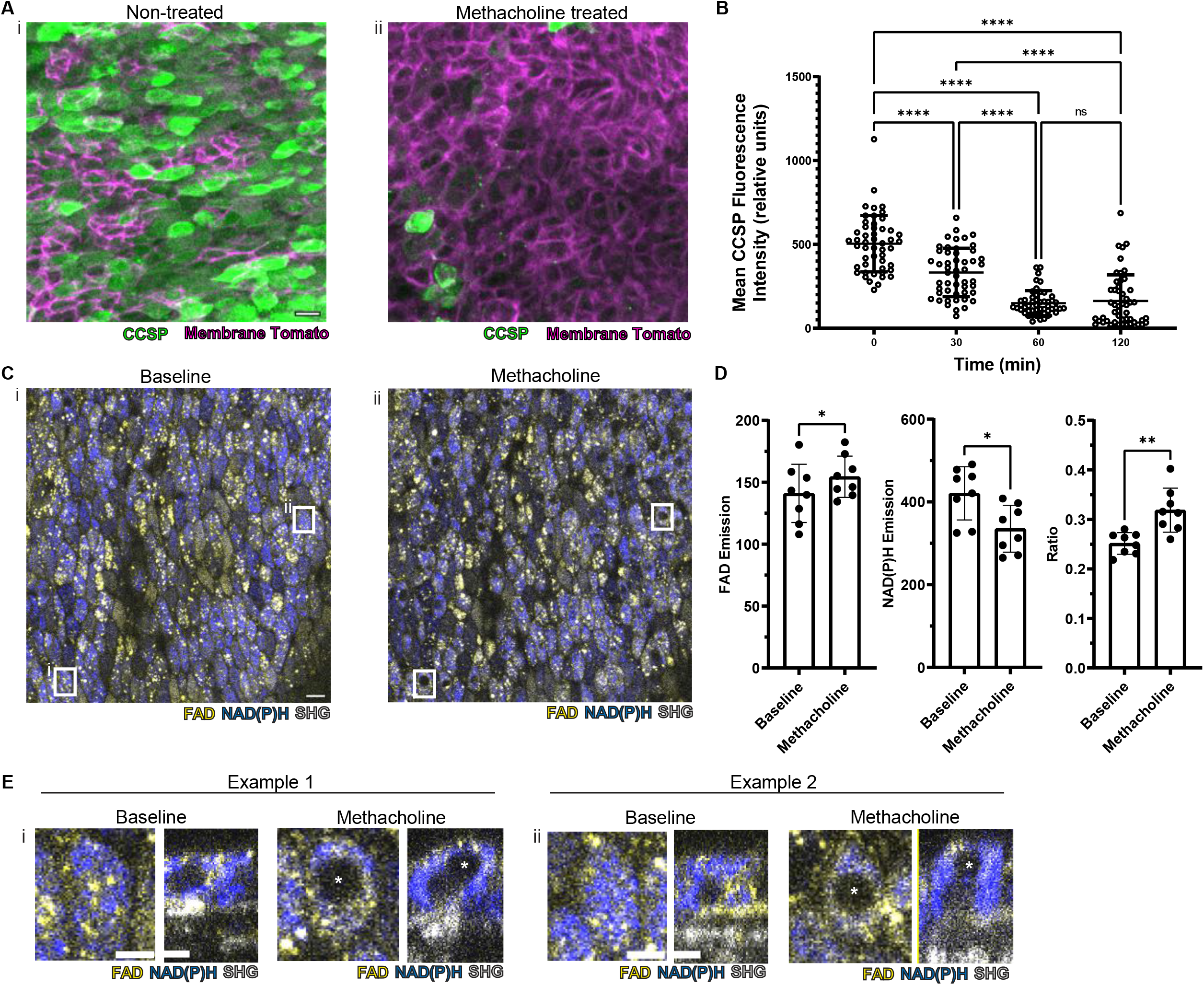
Autofluorescence imaging reliably identifies secretory cells while conventional CCSP staining is lost with methacholine stimulation. (A) Murine airway epithelial cells expressing membrane tomato under ROSA promoter. CCSP staining (green) using identical imaging parameters performed on two adjacent halves of same trachea (i) treated with vehicle vs (ii) 100uM methacholine showing decreased CCSP staining (green) after methacholine stimulation. Cell membrane denoted by membrane tomato (magenta). (B) Fluorescence intensity of CCSP staining over time with methacholine stimulation imaged with same parameters. Each cell is one dot. ****P<0.0001 by ANOVA. (C) Autofluorescence imaging (NADH blue, FAD yellow, SHG gray) of trachea at baseline (i) and after methacholine stimulation (ii). Zoom of white boxed region seen in (E). (D) Quantification of NADH, FAD, and fluorescence ratio (FAD/FAD+NAD(P)H) at baseline and after methacholine stimulation at single cell resolution (each dot is a cell). (E) White boxed regions from (C) showing the formation of non-fluorescent void after methacholine stimulation in XY view and cross-sectional XZ view. Non-fluorescent region in baseline is nucleus. Asterix denotes new non-fluorescent “void” after methacholine stimulation. Scale bars = 10 μm in A, C and 5 μm in E.

Identifying secretory cells using pre-stimulation autofluorescence intensity ratio, we found that methacholine stimulation does indeed alter NAD(P)H and FAD fluorescence intensities, however, the autofluorescence intensity ratio of secretory cells were still distinguishable from the surrounding epithelium cells. (Figure 4C-D). Thus, using this parameter, we were able to follow secretory cell identity after stimulation and were able to assess at single cell resolution, the response of a given secretory cell in real time.

To our surprise, we found that methacholine stimulation resulted in the development of non-fluorescent spherical “voids” within secretory cells (Figure 4C boxes, further magnification of these boxes in Figure 4E). We hypothesized that these novel structures were caused by an uptake of luminal extracellular contents. Indeed, such a process has been demonstrated in intestinal secretory/goblet cells (Gustafsson et al., 2021; Gustafsson & Johansson, 2022; McDole et al., 2012; Noah et al., 2019). To definitely evaluate our hypothesis concerning the origin of the spherical autofluorescence “voids” in our system, we added 10kd FITC-dextran to the extracellular fluid. Upon stimulation with methacholine, secretory cells showed uptake of FITC-dextran, distinct from the nucleus (Figure 5A). Since autofluorescence imaging alone cannot be used to establish the presence of a membrane, we resorted to employing epithelia expressing tdTomato on the membrane to see if the voids were membrane bound. Indeed, we found uptake of FITC-dextran into membrane encapsulated structures delineated by membrane tomato (Figure 5B). In order to visualize this process on a faster time scale, we utilized airway liquid interface cultures (ALI), which are significantly thinner than the native trachea allowing fast imaging across the Z axis. Using real time imaging, we captured the fine dynamics of the formation of the intracellular membrane bound structures that housed previously luminal contents. Still images demonstrate uptake of FITC-dextran into the cell from the apical surface, further accumulation of FITC-dextran into the cell, and a major secretion event (Figure 5C and Figure 5 Supplement Video). This process superficially bears a striking resemblance to the formation of intestinal goblet cell associated antigen passages (GAPs) in which an endocytic process in intestinal goblet cells is linked to mucin secretion (Gustafsson et al., 2021). We therefore tentatively refer to these structures as airway secretory cell associated antigen passages (SAPs) to indicate a potential similarity to their intestinal counterparts.

**Figure 5.**
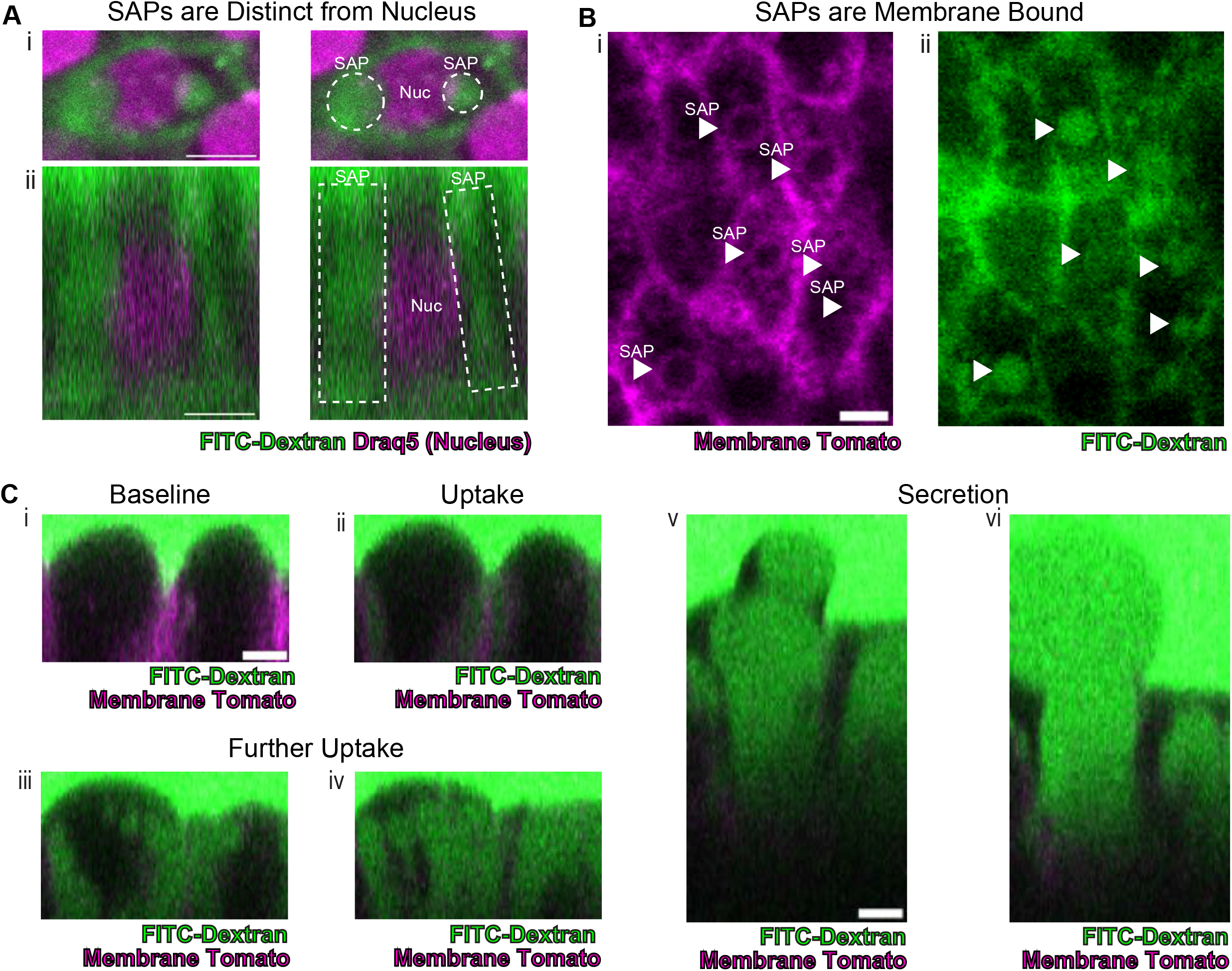
Secretory cells uptake luminal contents revealing secretory cell associated antigen passages (SAPs). (A) Secretory cell with nuclear stain Draq5 (magenta), demonstrating uptake of FITC-dextran (10 kD, green) is distinct from nucleus. (i) is xy view and (ii) is cross-sectional xz view. Dotted lines outlining “SAP” seen in secretory cell. Scale bars = 5 μm. After methacholine stimulation. (B) Trachea with (i) membrane tomato (magenta) and (ii) 10 kD FITC-dextran demonstrating methacholine stimulated uptake of extracellular dextran into membrane bound regions showing that SAPs (arrowheads) are membrane bound structures. Scale bars = 5 μm. (C) Selected images from Figure 5 Supplemental Video demonstrating uptake of luminal FITC-dextran and secretion of cell contents. Epithelium has membrane tomato (magenta) and FITC-dextran (green). (i) baseline image prior to methacholine stimulation (1 minute after methacholine stimulation). (ii) Uptake of FITC-dextran and formation of SAPs (9 minutes after methacholine stimulation). (iii-iv) Further accumulation of FITC-dextran (31 minutes and 40 minutes after methacholine stimulation). (v-vi) secretion of cellular contents (53 minutes and 60 minutes after methacholine stimulation). Scale bars = 5 μm.

**Figure 5 Supplement Video.** Murine Airway Liquid Interface culture (ALI) of airway epithelial cells labeled with membrane tomato (magenta) with surrounding environment containing FITC-dextran (green). Movie of SAP formation, demonstrating uptake of dextran into epithelial cell and secretion from the same cell. 4 frames per second. Imaging in the XZ plane. Scale bar = 5 μm.

## Discussion

We report a label free imaging methodology that can be applied to murine tracheal explants for cell type specific identification in live and unmanipulated tissues. We also show that autofluorescence coupled to morphology allows one to distinguish all the known airway epithelial cell types without the need for genetic labels.

Most importantly, we show that real time dynamic assessments of airway physiology are possible at a single cell resolution. Unexpectedly, cell identity can be established in a way that is sometimes more accurate than destructive immunofluorescent approaches. This may be of particular relevance to disease states, since it has been noted that markers of cell type such as the secretory cell marker CCSP can be significantly reduced in disease samples as in the case of COPD patient samples (Pilette et al., 2012). Similarly, CCSP positive cells have been shown to decrease in asthma (Shijubo et al., 2012) and in smokers in general (Shijubo et al., 1997). We speculate that secretory cells may persist in these pathologic states, but that they escape detection due to the loss of CCSP signal. Our label free imaging methodology is unaffected by such disease-relevant alterations in the expression of functional markers of epithelial cell type.

Finally, our identification of “secretory cell associated antigen passages” (SAPs) suggests that this new technology can be used to define a myriad of new real time physiologic processes that can then be subjected to deep mechanistic scrutiny.

Due to the simplicity of the explant model and the stability of this imaging technique over days of culture, this methodology could eventually be utilized to assess human airway surgical specimens and bronchoscopically obtained biopsies. If automated machine learning could be deployed, these techniques may eventually be useful for the real time assessment of disease-specific physiologic parameters and the real time assessment of the efficacy of therapeutic interventions aimed at restoring normal physiologic function.

## Materials and Methods

### Mouse Models

C57BL/6J mice (stock no. 000664) and MT-MG (stock no. 007676), were purchased from Jackson laboratory.

Mice were maintained in accordance with the Association for Assessment and Accreditation of Laboratory Animal Care-accredited animal facility at the Massachusetts General Hospital. All procedures were performed with Institutional Animal Care and Use Committee (IACUC)- approved protocols. All mice were housed in an environment with controlled temperature and humidity, on 12-hour light:dark cycles, and fed with regular rodent’s chow.

### Mouse Tracheal Explant Model

As demonstrated in Figure 1 Supplement 1A, mice were euthanized as per MGH IACUC protocols and trachea were dissected, cleared of connective tissue and opened longitudinally and bisected. The dissected trachea was then sutured onto a silicone O-ring and placed onto a custom made insert. The custom made insert was assembled using an inverted 6.5 mm Transwell®, 0.4 μm Pore Polyester Membrane Insert fixed to a custom 3D printed insert (Supplemental Material) with Sylgard 184 silicone elastomer (EMS Catalog #24236-10).

Samples were incubated with Gibco DMEM/F12 (Fischer 11320033). For experiments across multiple days, the insert was filled with media such that the meniscus was located below the trachea to prevent explant submersion. Primocin (InVivoGen) was added to the media to prevent contamination.

### Two-photon Microscopy

Tracheal explants were fixed to inserts as described above. An Olympus FVMPE-RS multiphoton laser scanning microscope equipped with a MaiTai HPDS-O IR pulsed laser (900 nm for FAD and SHG) and INSIGHT X3-OL IR pulsed laser (730 nm for NAD(P)H), using a 25X water immersion lens (NA 1.05) (XLPN25SWMP2) for imaging. Media was used as the submersion droplet. Configuration of microscope setups is seen in Figure 1 Supplement 1B. Additionally, a 2x optical zoom with 1024×1024 scan size was used. A full thickness Z section was imaged from above the tracheal surface through to the sub epithelium, with a 0.5μm step size. Multiple regions on the same tracheal explant were imaged. Each region was “registered” using SHG features of the subepithelial collagen/cartilage/vasculature as fiduciary marks. The insert was placed in a physiological live imaging chamber (CO_2_ and temperature-controlled, TokaiHit) at 37°C and 5% CO_2_.

### Mitochondrial Poisons

Tracheal explants were incubated with DMEM/F-12 Media with Primocin (InVivoGen) and 15 mM HEPES. After baseline imaging, samples were incubated with complex 1 inhibitor - Antimycin A (10uM) and complex III inhibitor - Rotenone (1uM) for 10 minutes mixed in the above media. Imaging was conducted as described. Similarly, after baseline imaging, tracheal explants were incubated with media containing mitochondrial uncoupler, Carbonyl cyanide p-trifluoro-methoxyphenyl hydrazone (FCCP, 1uM).

### Immunofluorescence staining of Tracheal Explants

Tracheal explants were secured onto the insert setup as described above. Tracheas were rinsed in PBS and subsequently fixed in 4% PFA in PBS for 20 minutes at room temperature. A glass slide was placed a few millimeters above the tissue to form a column of liquid. Explants were subsequently washed with PBS three times for a total of 1 hour. The explant were then permeabilized in PBS-0.3% TritonX-100 (PBST) for 30 minutes in a similar manner. The sample was then stained with primary antibody at 37°C for 1 hour, diluted in 1%BSA-0.3%PBST. The following antibodies and dilutions were used:

| Antibody | Manufacturer | Dilution |
| --- | --- | --- |
| mouse anti-tubulin, acetylated | Sigma T-6793 | 1:500 |
| rabbit anti-BSND | Abcam ab196017 | 1:150 |
| goat anti-GNAT3 | Novus NBP1-20926 | 1:100 |
| rabbit anti-KRT13 | Abcam ab92551 | 1:100 |
| UEA1-dylight 649 | Vector Laboratories DL-1068-1 | 1:100 |
| goat anti-CC10 | Santa Cruz SC-9772 | 1:1000 |
| rabbit anti-UCHL1 | Proteintech 14730-1-AP | 1:250 |
| chicken anti-KRT5 | Biolegend 905901 | 1:200 |
| Draq5 | Abcam ab108410 | 1:5000 |

Tracheal explants were then washed with PBS for 1 hour and stained with secondary antibody at 37°C for 1 hour. Various secondary antibodies (Jackson ImmunoResearch) were used. The samples were then imaged using 2-photon microscopy as above. The Olympus FVMPE-RS multiphoton laser scanning microscope was equipped with a MaiTai HPDS-O IR pulsed laser (900 nm for AF488, AF647, and SHG) and an INSIGHT X3-OL IR pulsed laser (1050 nm for AF594, Draq5), using a 25X water immersion lens (NA 1.05). Samples were registered to the identical location of autofluorescence microscopy based on SHG features of the subepithelial collagen/cartilage/vasculature as fiduciary marks.

### Cell Autofluorescence and Morphological Analysis

Autofluorescence images were obtained as described above. In order to normalize the Z-axis to the basement membrane (SHG), or “flatten” the image, a matlab algorithm (Appendix 1) was utilized. With the airway epithelial cells in a similar plane, the autofluorescence intensity was measured at the epithelial surface in the NAD(P)H and FAD channels using Image J. The morphological measurements were conducted as described in Figure 3 Supplement 1.

### Dimensionality reduction and visualization

UMAP was calculated using the package in Python with the following parameters: random_state = 42; spread = 3; min;_dist =0.1. A total of 206 cells were analyzed. Data visualization was implemented using the matplotlib package in Python. All the codes are provided in the supplemental jupyter notebook file (Appendix 2).

### Machine learning algorithms for classifying cell types

K nearest neighbor, multinominal logistic regression, and XGBoost algorithms were used to classify cell types. The Scikit-learn package was used to implement k nearest neighbor and multinominal logistic regression algorithms and XGBoost package was used to implement XGBoost algorithm. All the codes are provided in the supplemental jupyter notebook file. The following features were used to train the algorithms: 1. FAD; 2. NADH; 3. FAD/NADH ratio; 4. Nuclear position; 5. X-length; 6. Y-length; 7. Z-length; 8. XY aspect ratio; 9. YZ aspect ratio; 10. Standard deviation of the fluorescence ratio above nucleus (Top std); 11. Standard deviation of fluorescence ratio below nucleus (Bot std); 12. Top std/Bot std ratio. 152 cells (75%) were used to train the model and 50 cells (25%) were used to test the models. For k nearest neighbor algorithm, k = 3 yields the highest accuracy and was therefore used to generate the confusion matrix. The Matthew’s Correlation Coefficient was calculated using the Scikit-learn package.

### Methacholine Treatment and CCSP Immunofluorescence staining

Tracheal explants from mice expressing membrane tomato under the ROSA promoter (stock no. 007676) were euthanized and dissected as described above. Tracheal samples were bisected as described with one half placed in DMEM/F12 media while the other half was placed into DMEM/F12 with 10uM of methacholine (dry powder dissolved in DMEM/F12, Sigma A2251). After 30 minutes, 1 hours, 1.5 hours, and 2 hours of incubation, the samples were fixed in 4% PFA in PBS for 20 minutes at room temperature with agitation. Samples were then washed with PBS three times for a total of 1 hour. Samples were permeabilized in PBS-0.3% Triton X-100 (PBST) for 30 minutes with gentle agitation. The samples were stained with CC10 primary antibody (aka Scgb1a1, 1:500; SC-9772, Santa Cruz) at 37°C for 2 hours, diluted in 1% BSA- 0.3% PBST. Samples were rinsed in PBS for 30 minutes and incubated into secondary antibody for 30 minutes with a subsequent 30 minutes PBS wash. Imaging was conducted on the Olympus FVMPE-RS multiphoton laser scanning microscope equipped with a MaiTai HPDS-O IR pulsed laser (900 nm for AF488) and an INSIGHT X3-OL IR pulsed laser (1050 nm for membrane tomato). Laser power was kept constant for all images and all tracheal samples were acquired using the same settings. ImageJ was used for quantifying fluorescence intensity on maximal projection images.

### FITC-Dextran Uptake Assays

Explants were prepared as described. If secretory cell identification was required, autofluorescence imaging was conducted as described. Explants were then incubated with 10 kd FITC-dextran (Sigma FD10S) diluted into DMEM/F12 media at a concentration of 1mg/ml with 10μM methacholine (dry powder dissolved in DMEM/F12, Sigma A2251). Imaging was conducted on the Olympus FVMPE-RS multiphoton laser scanning microscope equipped with a MaiTai HPDS-O IR pulsed laser (800 nm for FITC) and an INSIGHT X3-OL IR pulsed laser (1050 nm for Draq5 or membrane tomato) with 2x optical zoom, 1024×1024 scan size, and a step size of 0.5μm.

### Air Liquid Interface Model

Trachea cells were harvested and cultured as previously described (Mou et al., 2016). Briefly, cells were cultured and expanded in complete SAGM (small airway epithelial cell growth medium; Lonza, CC-3118) containing TGF-β/BMP4/WNT antagonist cocktails and 5 μM ROCK inhibitor Y-27632 (Selleckbio, S1049). To generate air-liquid interface (ALI) cultures, airway basal stem cells were seeded onto transwell membranes and differentiated using PneumaCult-ALI Medium (StemCell, 05001). Air-liquid cultures were fully differentiated after 14 days as demonstrated by the formation of air-liquid interface and the beating of cilia.

### Statistical Analysis

All analysis was performed using Graphpad Prism. For analysis of experimental data after treatment with mitochondrial poisons (Figure 1C-D), imaging over time (Figure 1E), and methacholine treatment (Figure 4D), a paired t test was used. For comparison of autofluorescence ratio across different cell types (Figure 2D), one-way ANOVA was utilized. For comparison of immunofluorescence staining with various durations of methacholine treatment (Figure 4B), a one-way ANOVA was used.

## Author Details, Contributions, and Competing Interests

Viral Shah (VS)

- Division of Pulmonary and Critical Care Medicine, Department of Medicine, Massachusetts General Hospital, Boston, MA, USA.
- Center for Regenerative Medicine, Massachusetts General Hospital, Boston, MA, USA.

**Contributions:** Conceptualization, Investigations, Methodology, Visualization, Analysis, Writing (original draft, review, editing)

**Competing Interests**: None.

Jue Hou (JH)

- Advanced Microscopy Program, Center for Systems Biology and Wellman Center for Photomedicine, Massachusetts General Hospital, Harvard Medical School, Boston, MA, USA.

**Contributions:** Conceptualization, Investigations, Methodology, Visualization, Analysis, Writing (review and editing)

**Competing Interests**: None.

Vladimir Vinarsky (VV)

- Center for Regenerative Medicine, Massachusetts General Hospital, Boston, MA, USA.

**Contributions:** Conceptualization, Investigations, Methodology

**Competing Interests**: None.

Jiajie Xu (JX)

- Center for Regenerative Medicine, Massachusetts General Hospital, Boston, MA, USA.

**Contributions:** Investigation, Methodology, Writing (review and editing)

**Competing Interests**: None.

Charles P Lin (CPL)

- Advanced Microscopy Program, Center for Systems Biology and Wellman Center for Photomedicine, Massachusetts General Hospital, Harvard Medical School, Boston, MA, 02114

**Contributions:** Conceptualization, Investigations, Methodology, Analysis, Writing (review and editing), Supervision, Resources, Funding acquisition

**Competing Interests**: None.

Jayaraj Rajagopal (JR)

- Division of Pulmonary and Critical Care Medicine, Department of Medicine, Massachusetts General Hospital, Boston, MA, USA.
- Center for Regenerative Medicine, Massachusetts General Hospital, Boston, MA, USA.
- Harvard Stem Cell Institute, Cambridge, MA, USA.
- Klarman Cell Observatory, Broad Institute of MIT and Harvard, Cambridge, MA, USA.

**Contributions**: Conceptualization, Investigations, Methodology, Analysis, Writing (review and editing), Supervision, Resources, Funding acquisition

## Acknowledgements

We would like to thank the members of the Rajagopal lab and Lin labs for their feedback. We would like to thank Manalee Surve for her discussions and input. This work was supported by NIH-NHLBI 5R01HL142559-04 (JR, CPL), RO1HL118185-08 (JR), 1R01HL157221-01A1 (JR), 5R01HL148351-03 (JR), Bernard and Mildred Kayden Endowed MGH Research Institute Chair (JR), and CFF 003338L121 (VS).

## Appendix 1

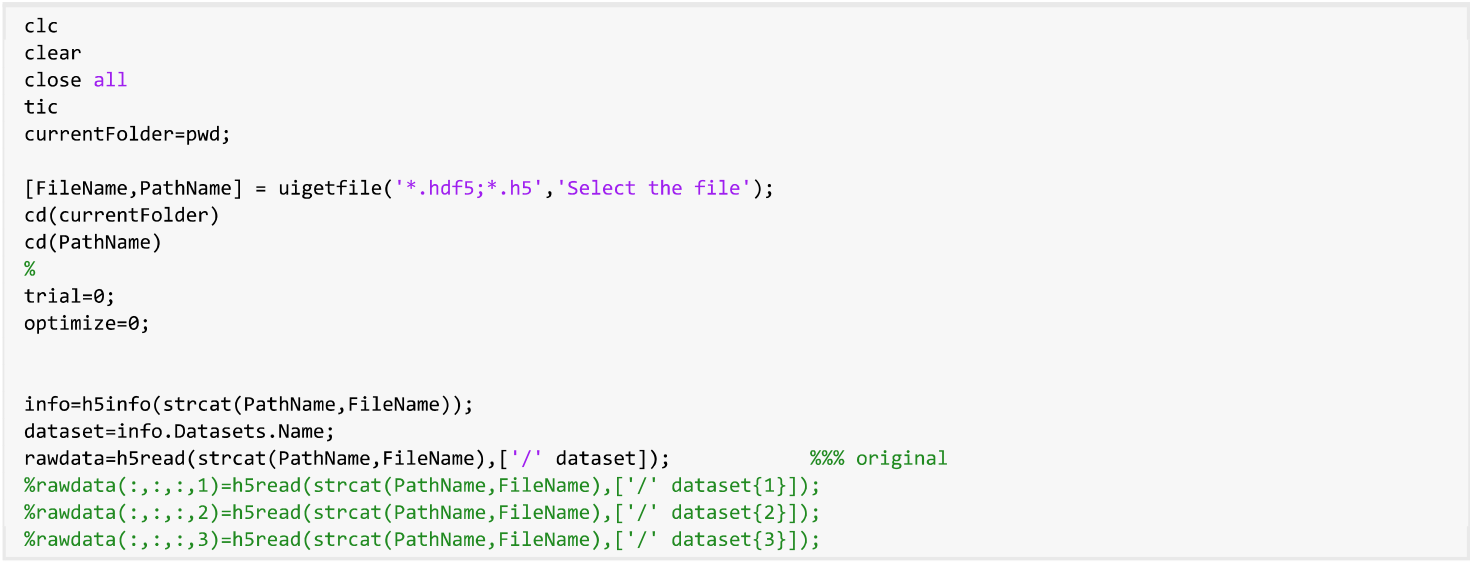

### select shg

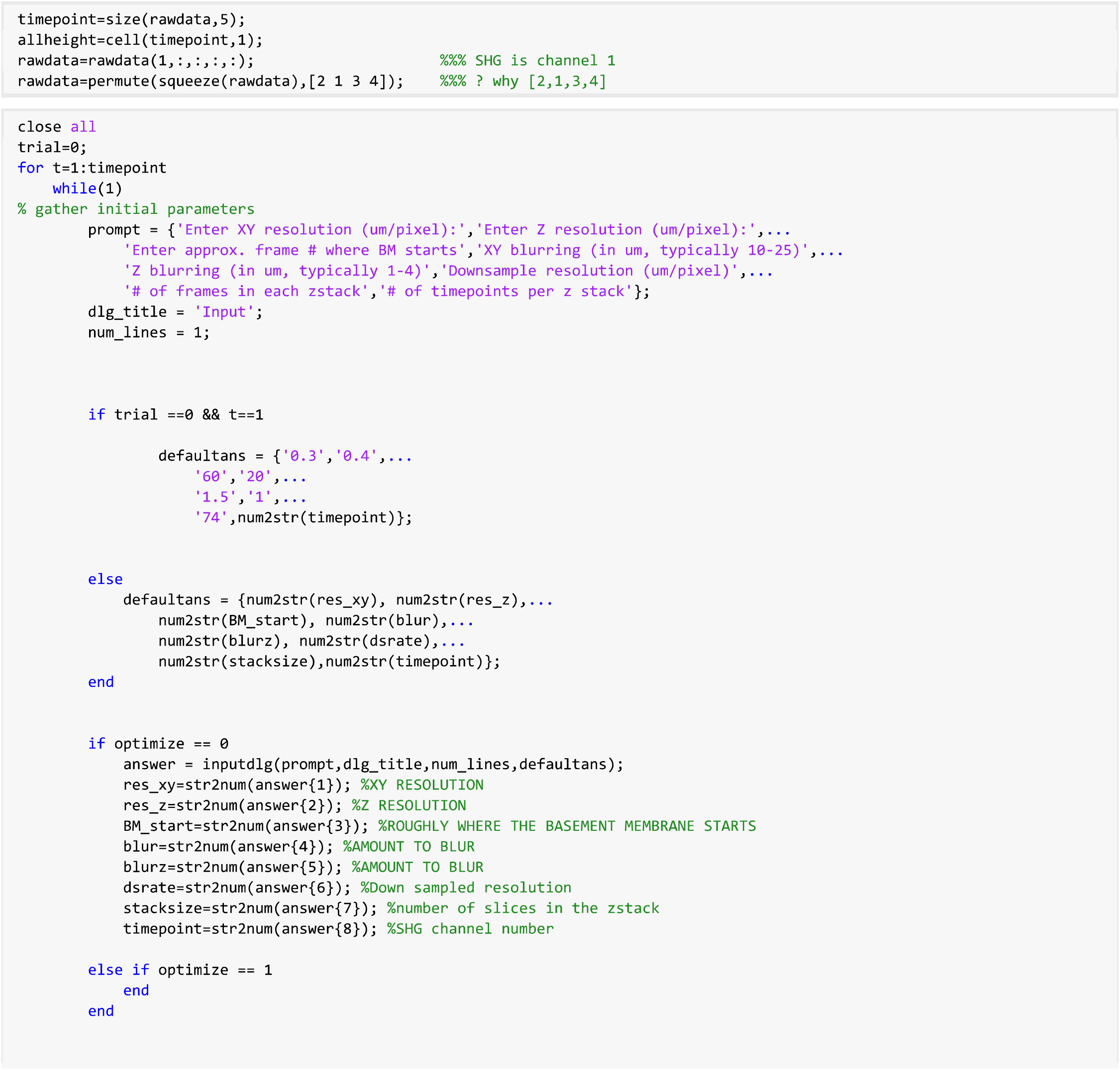

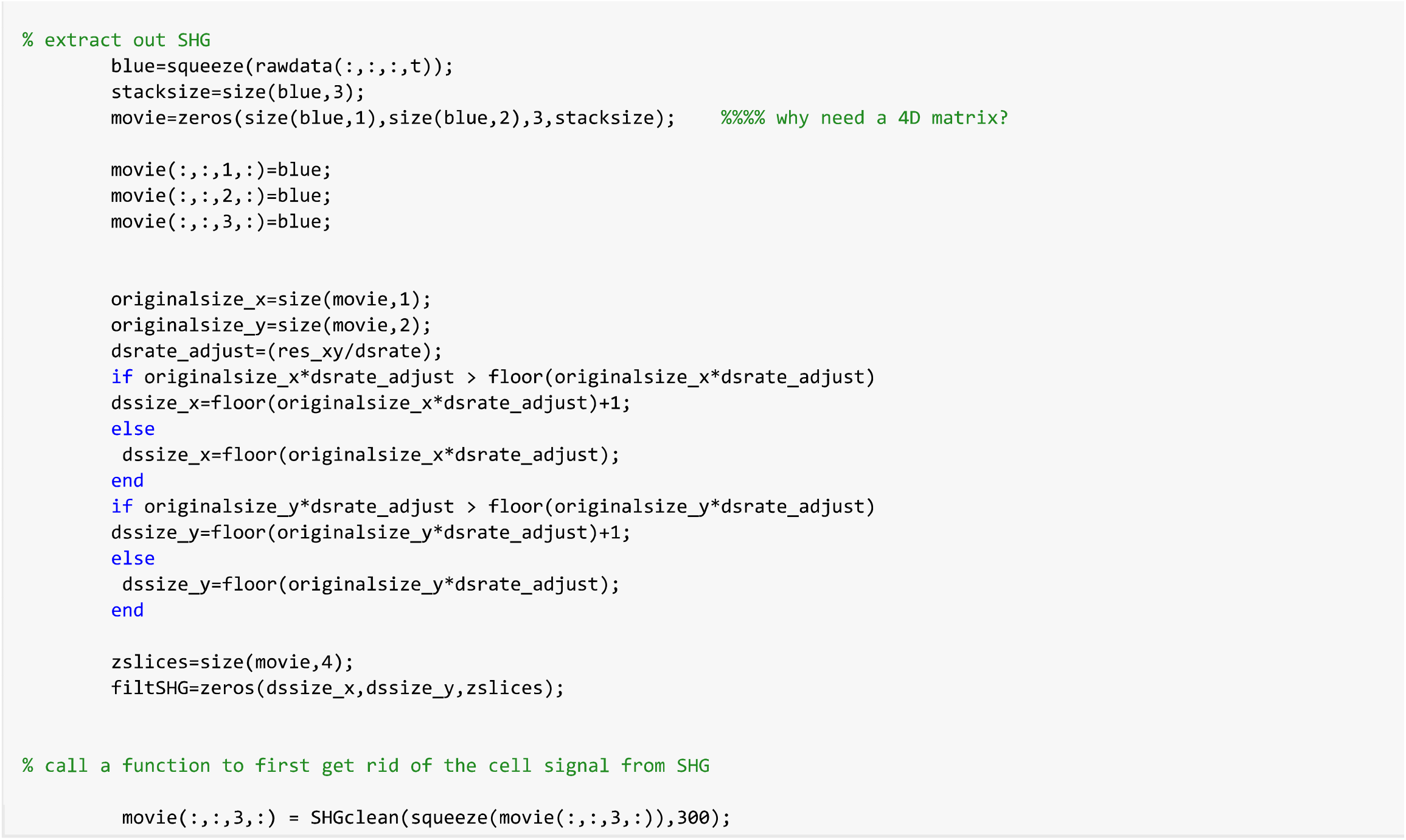

### blur the SHG

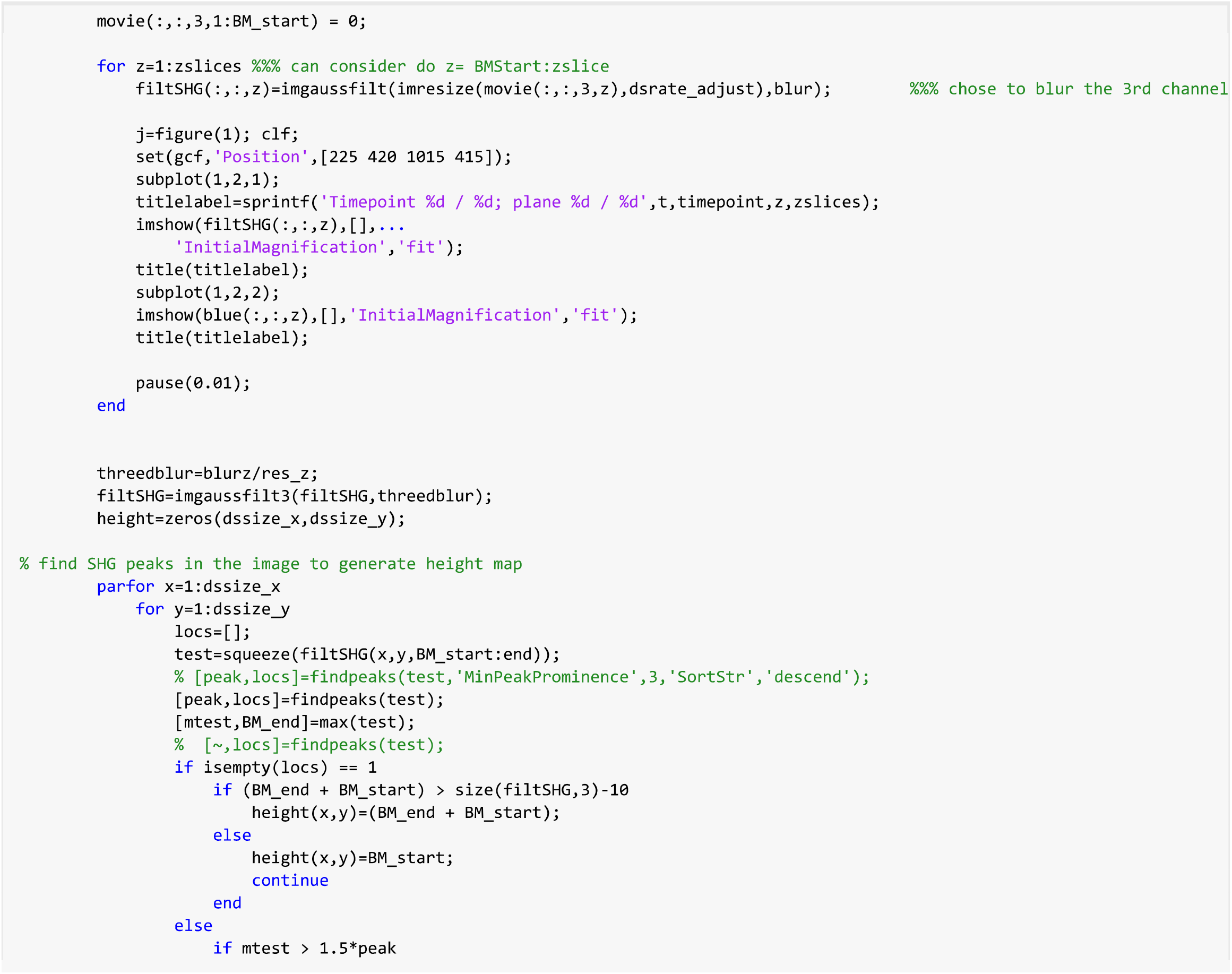

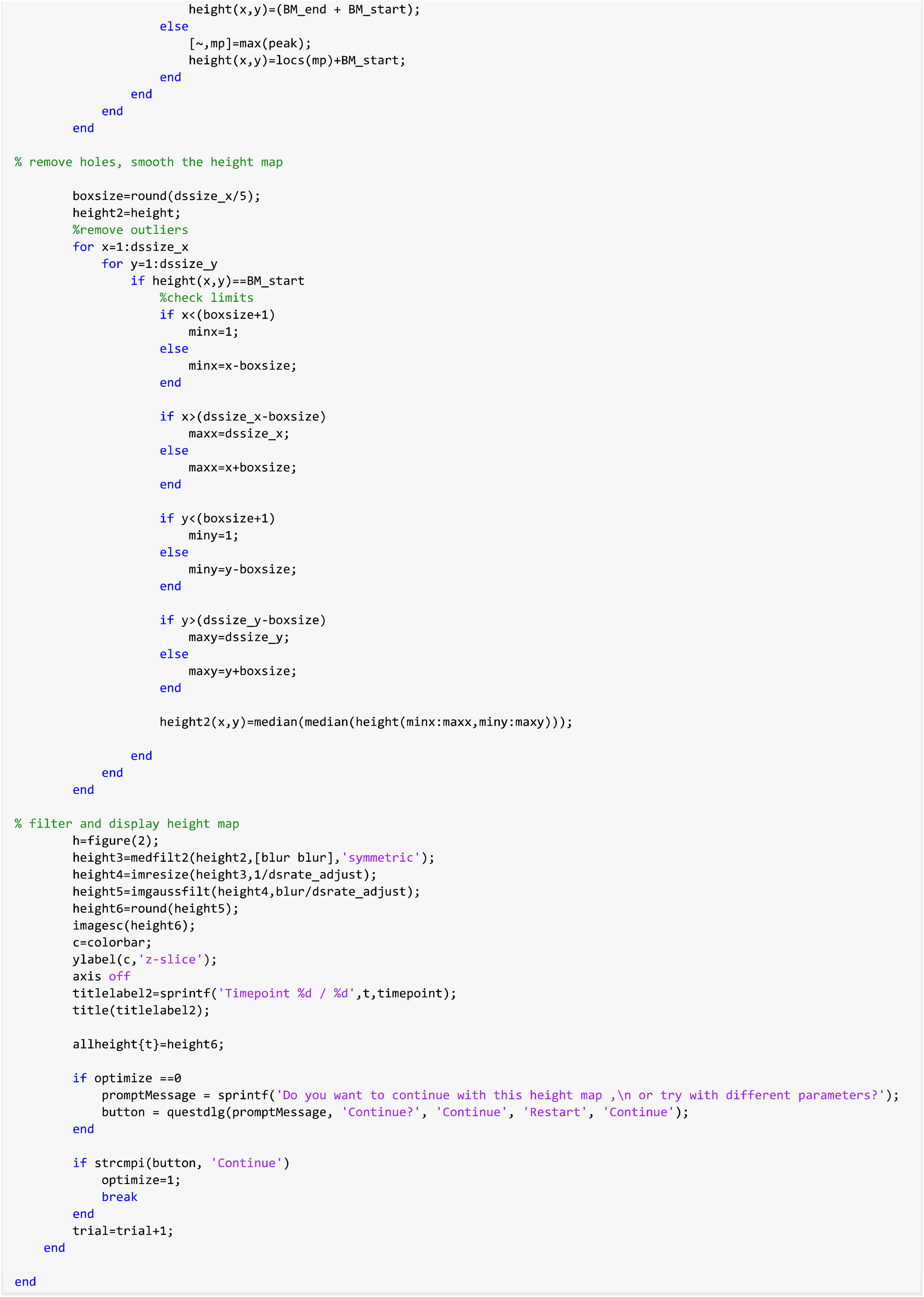

### plot curvature of z stack

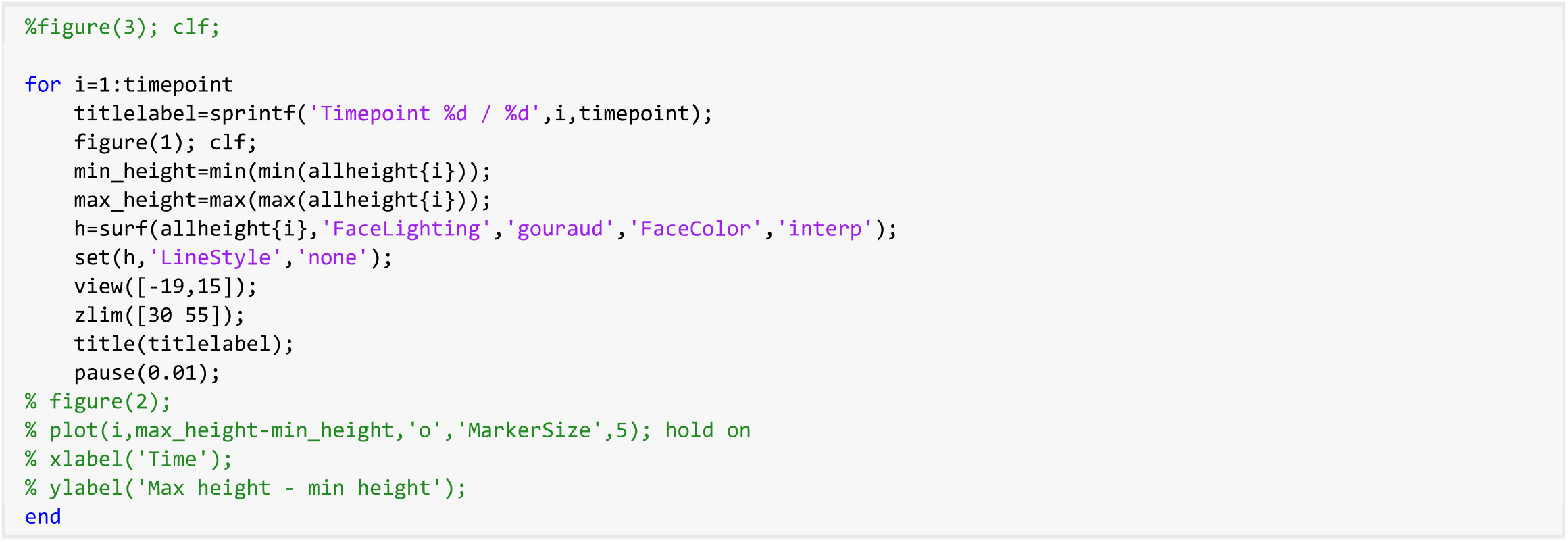

### IF HEIGHTMAP IS GOOD, SAVE allheight as allheight.mat %%%%%%%%%%%%

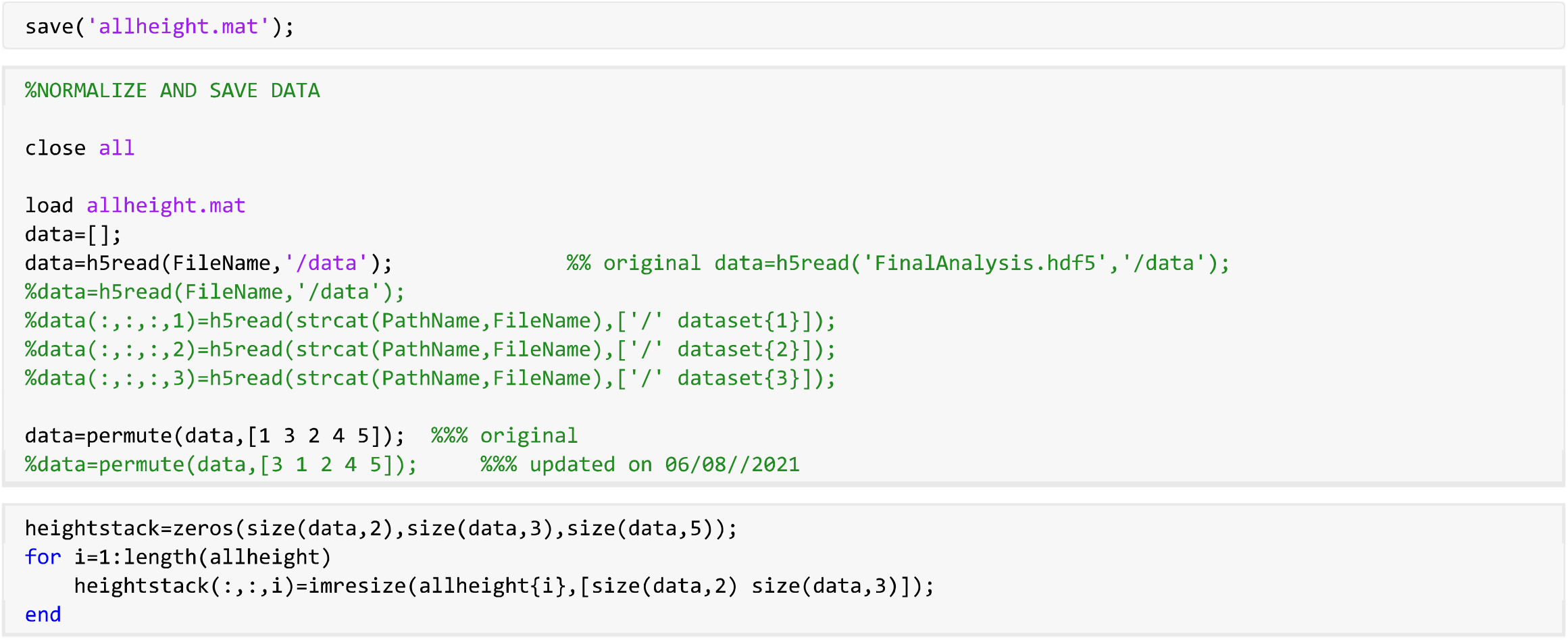

### norrmalize

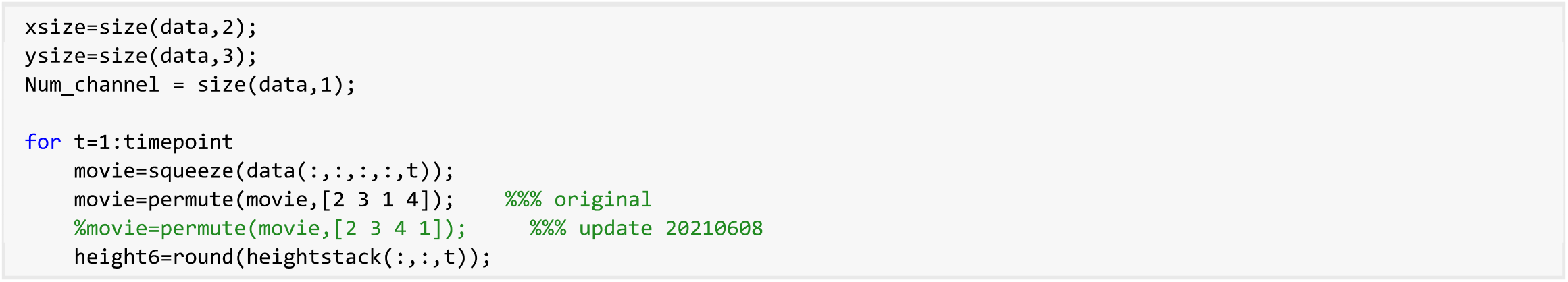

### limit max height to not exceed stack

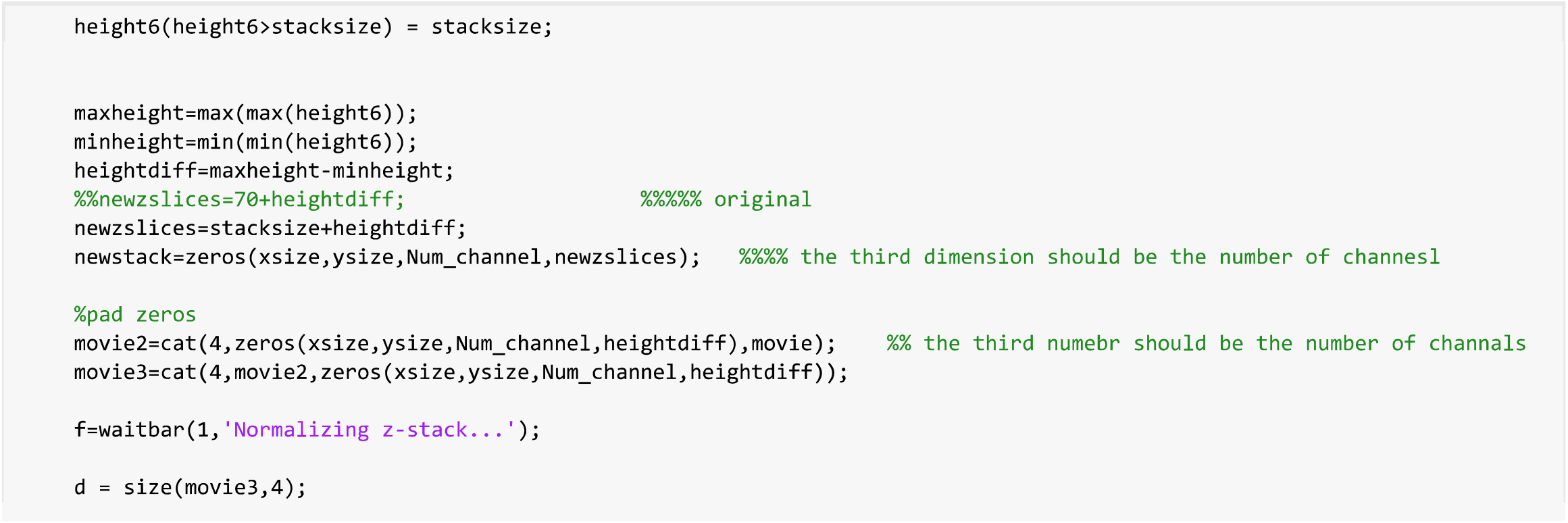

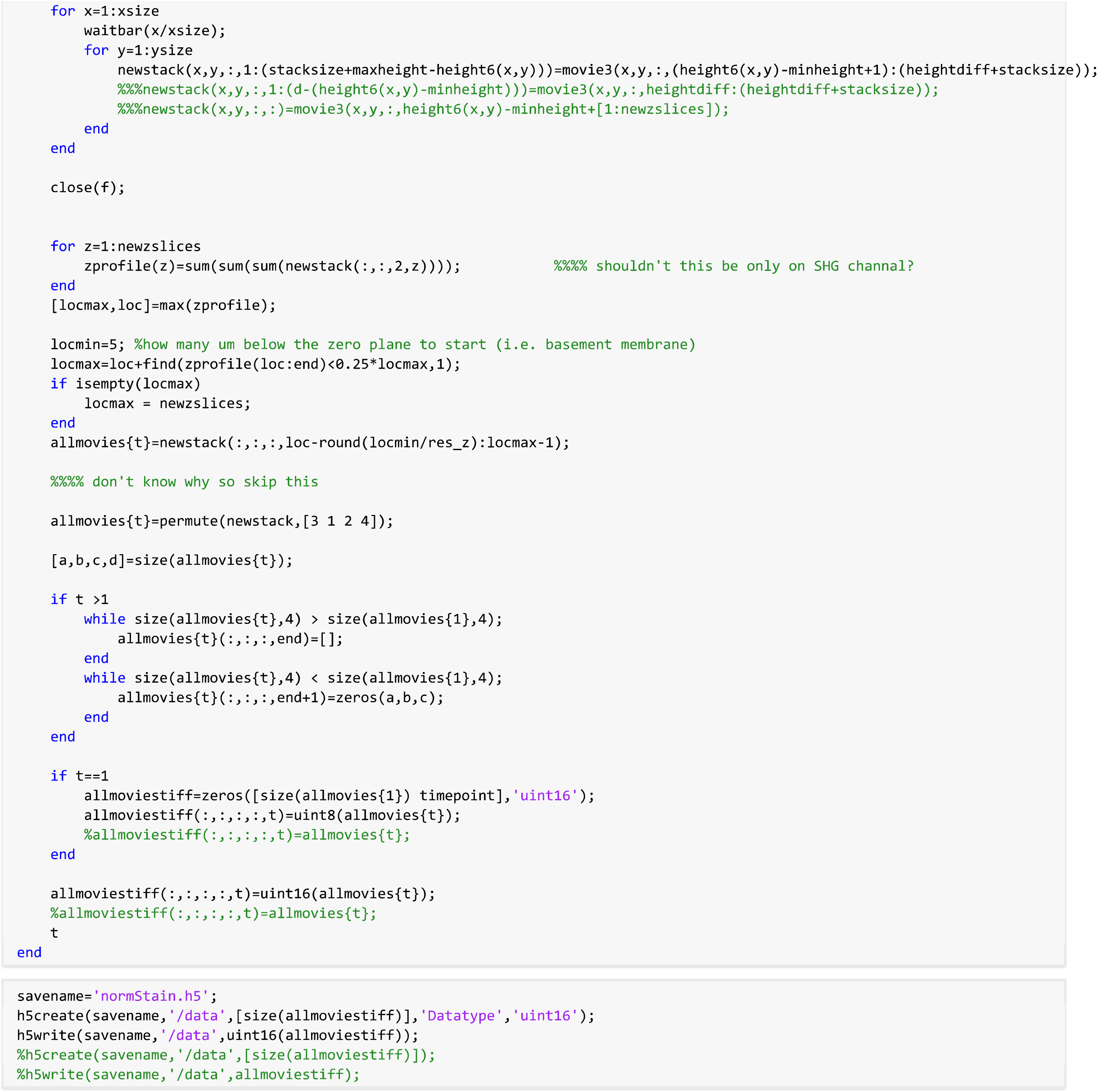

## Appendix 2

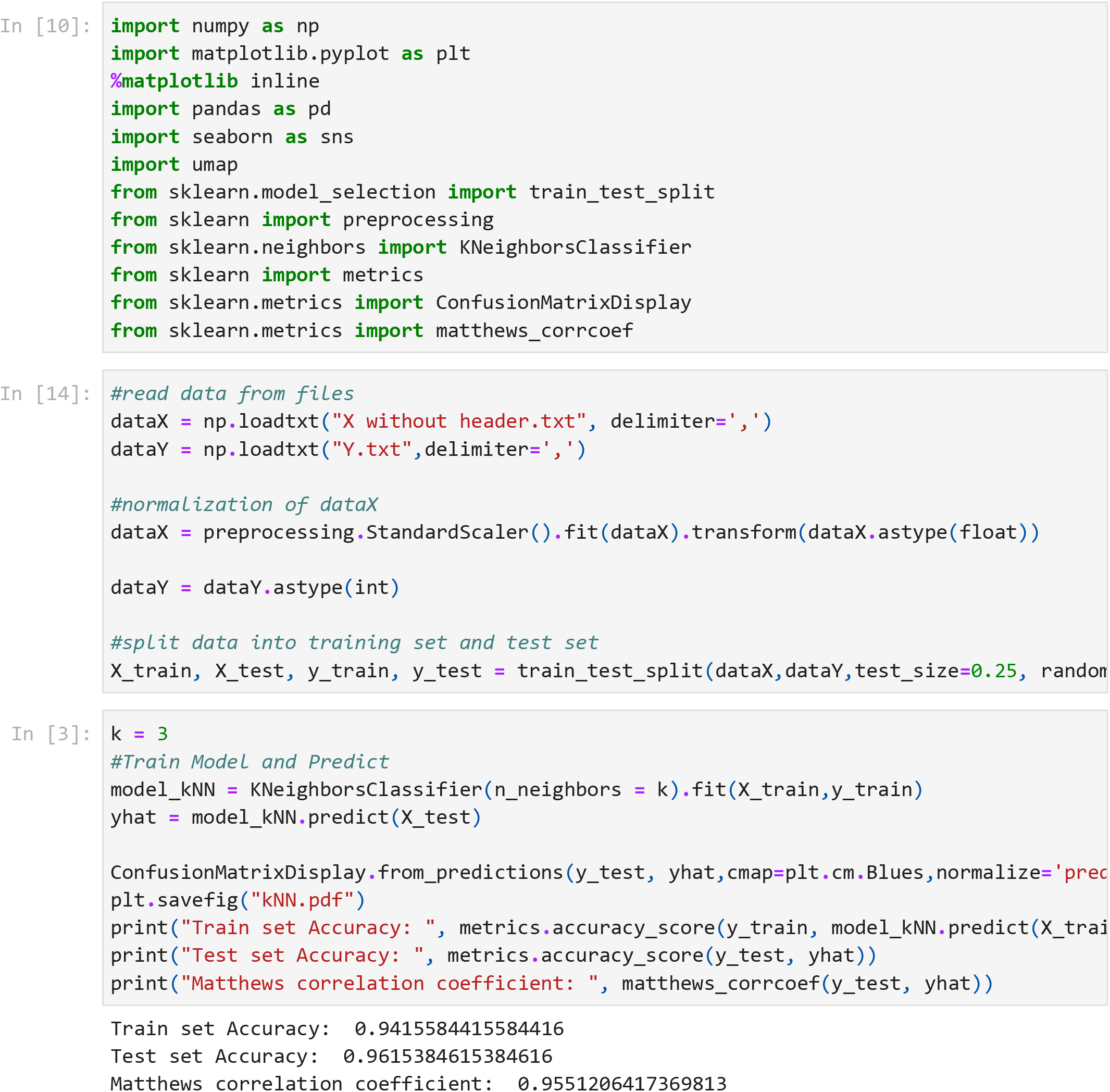

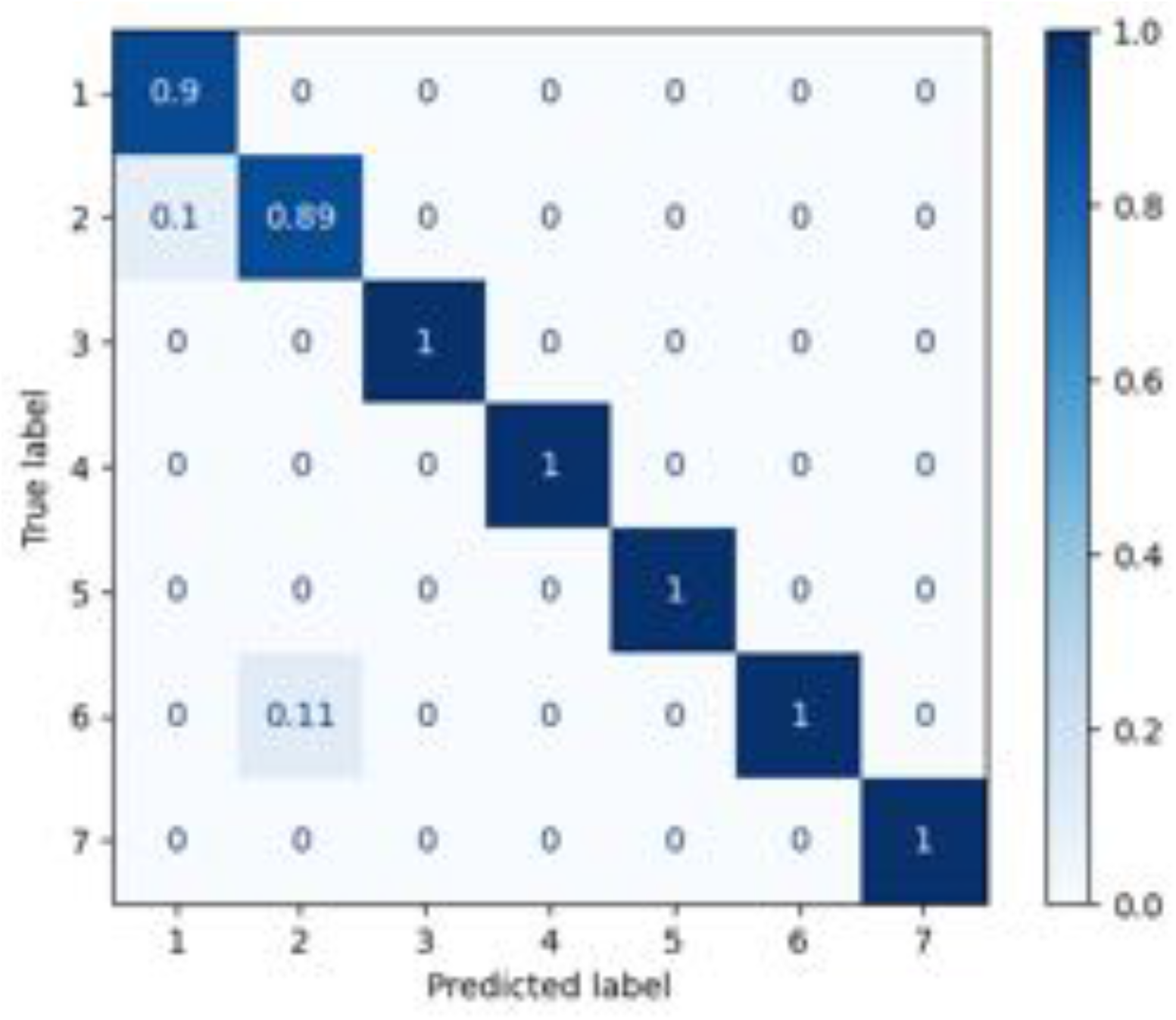

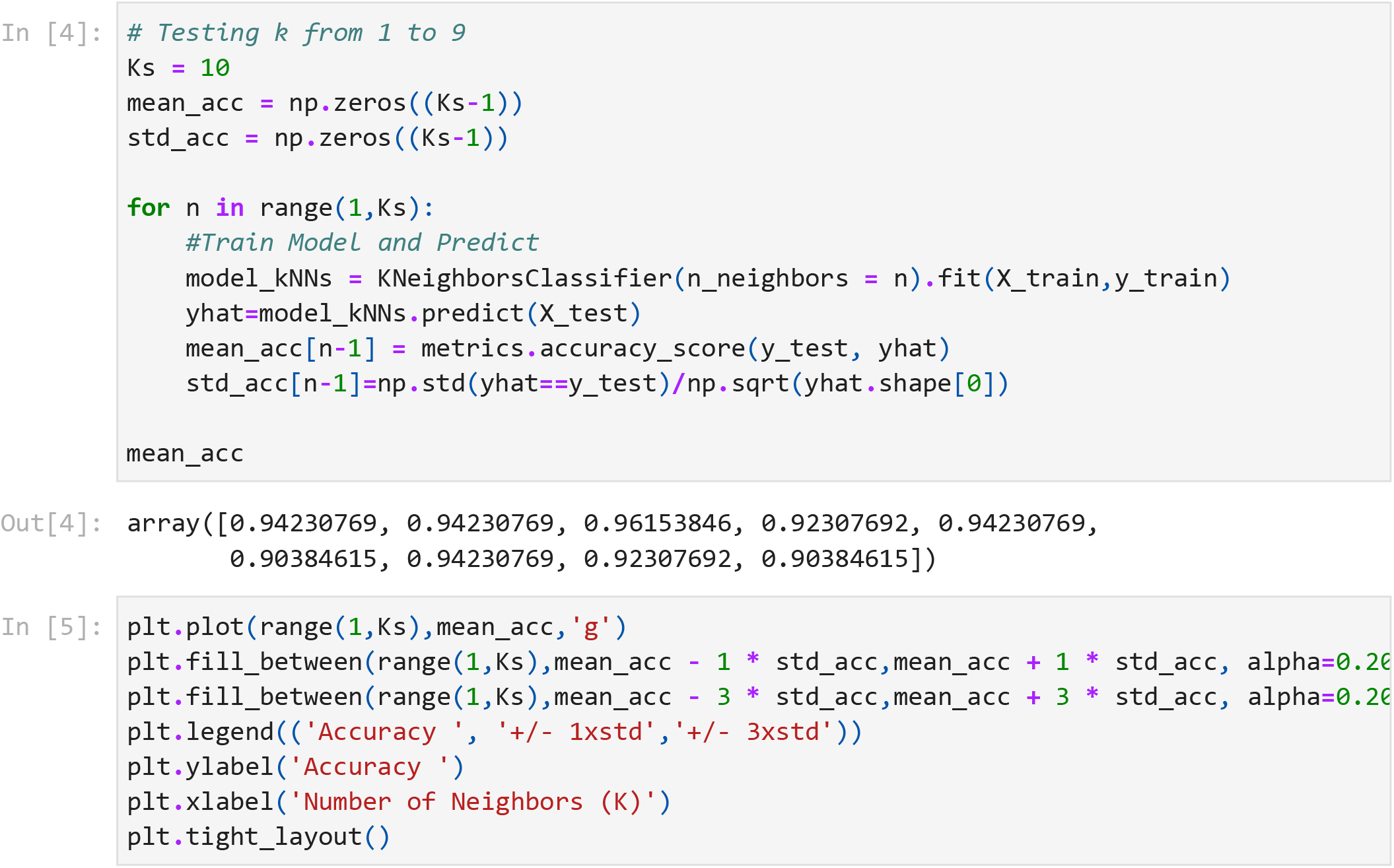

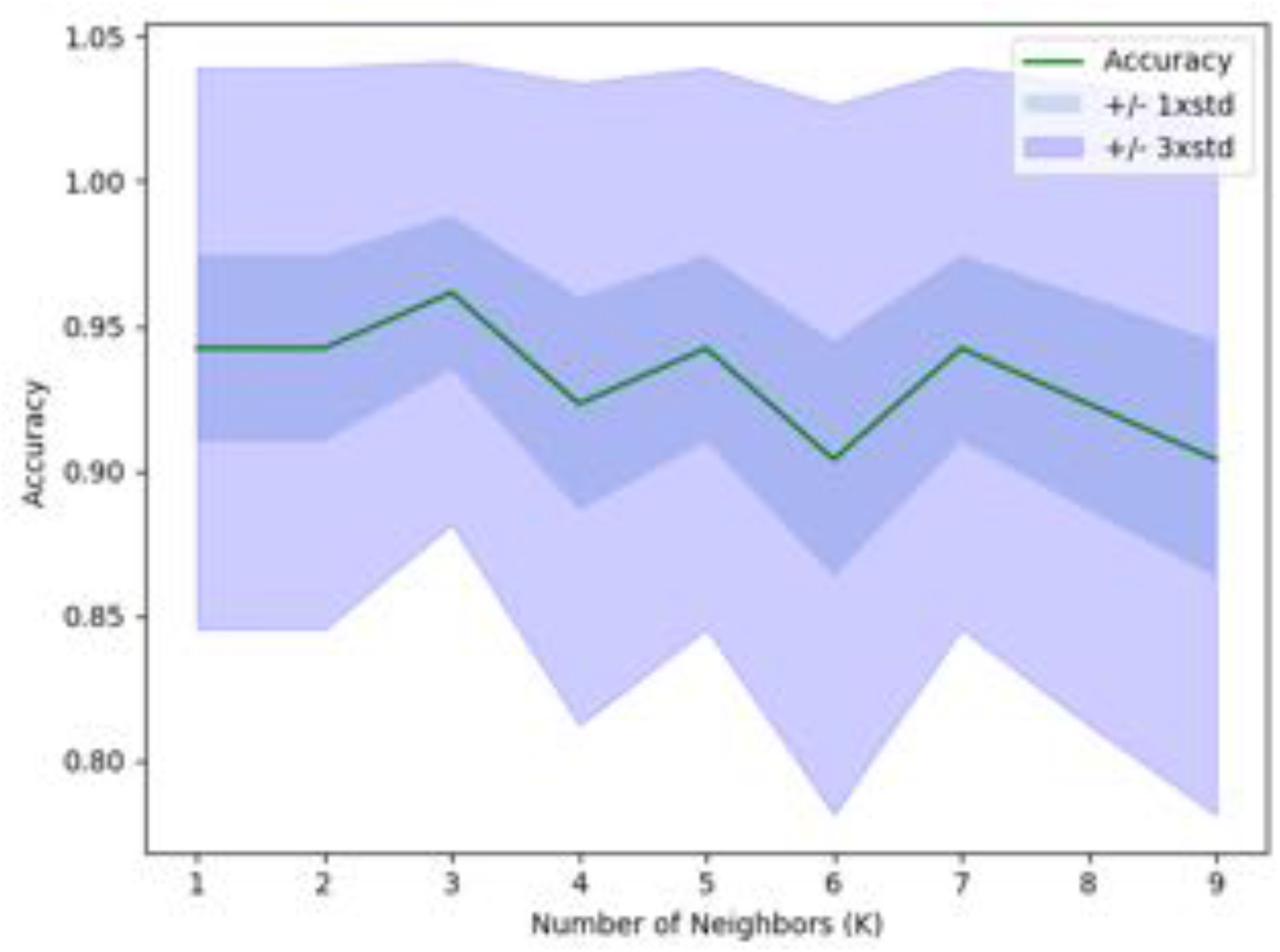

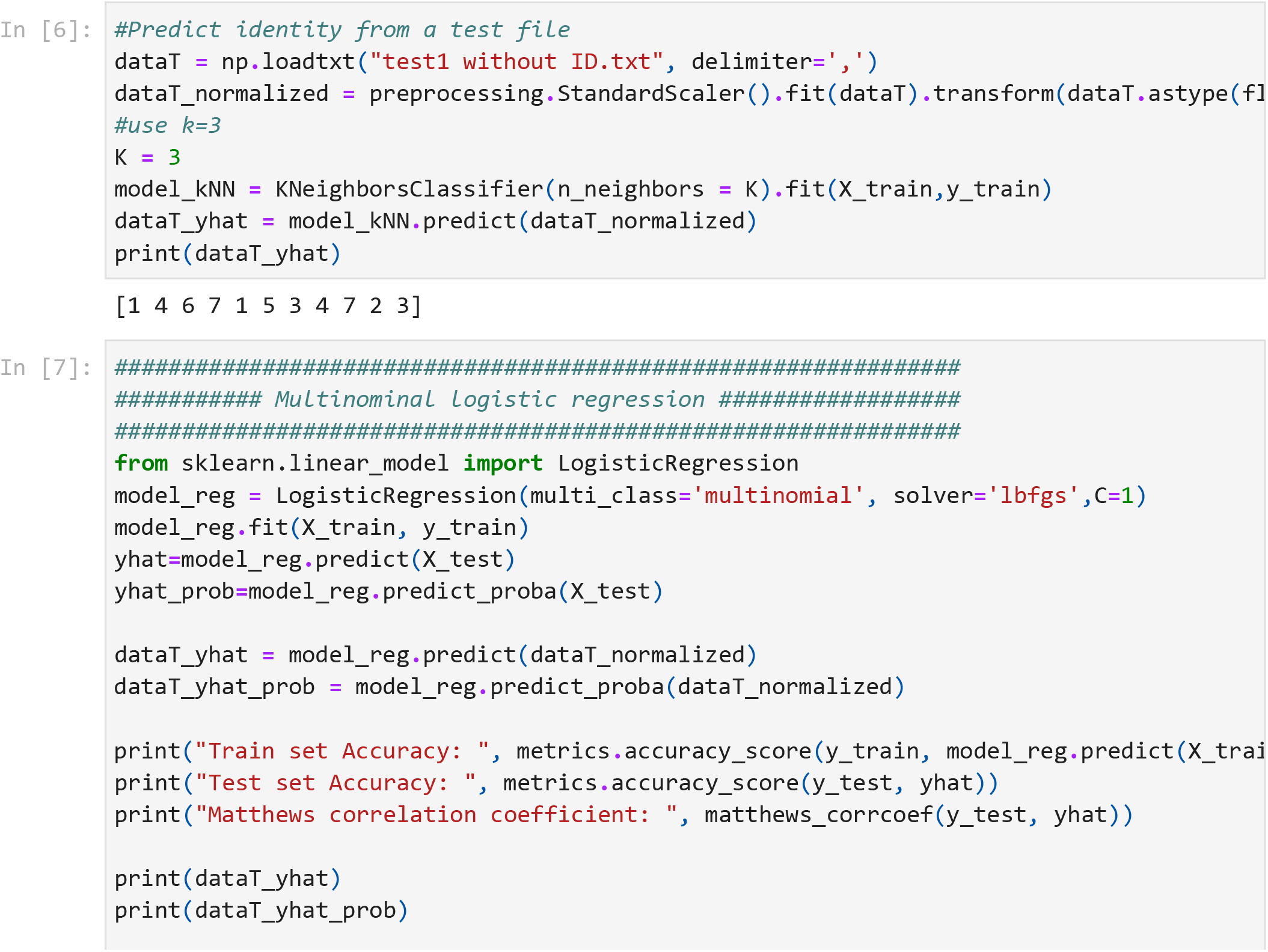

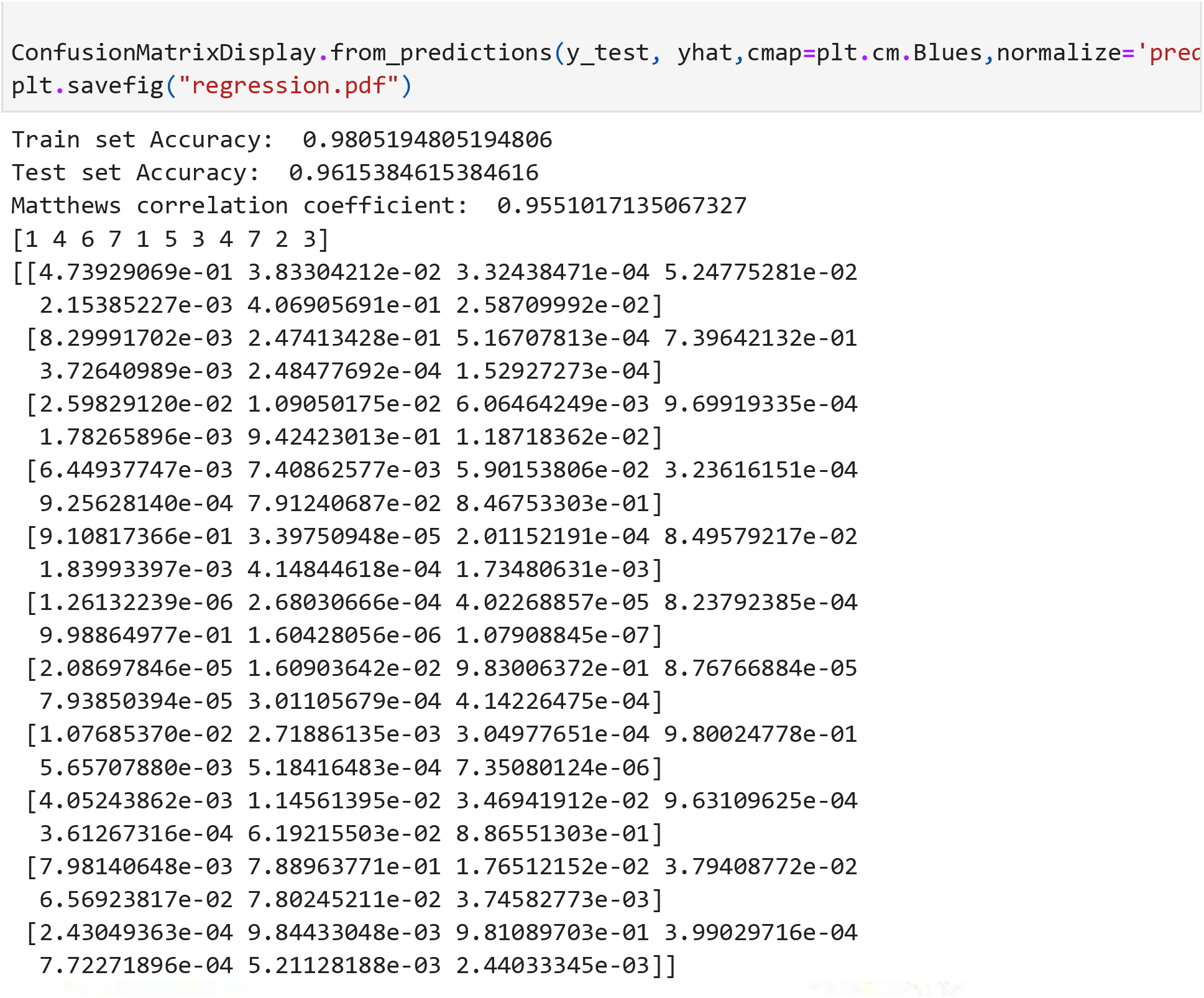

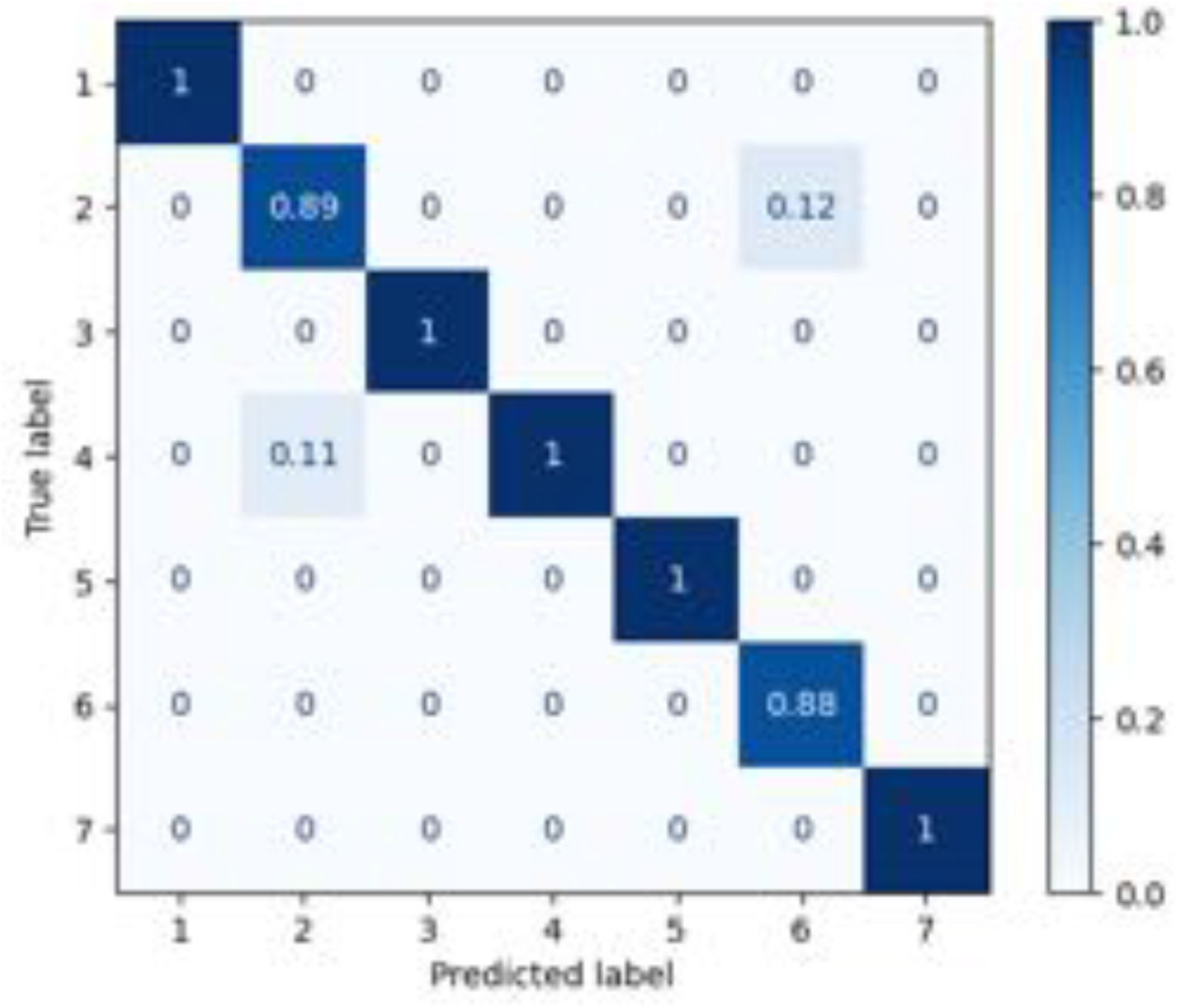

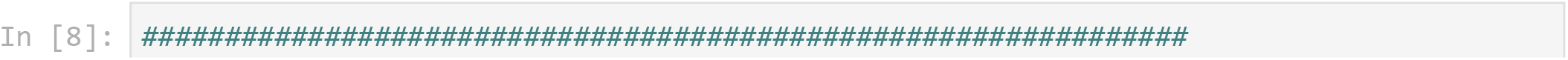

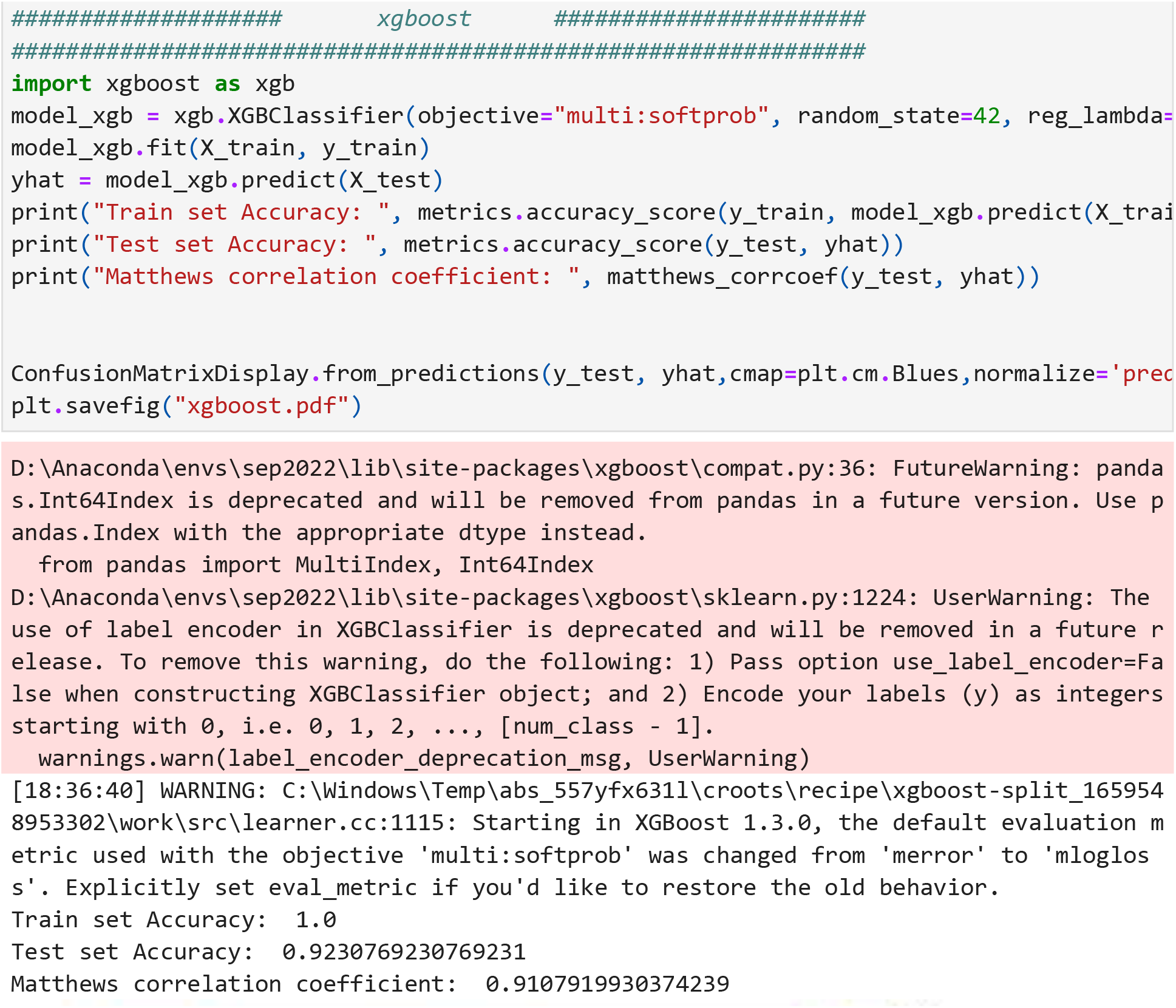

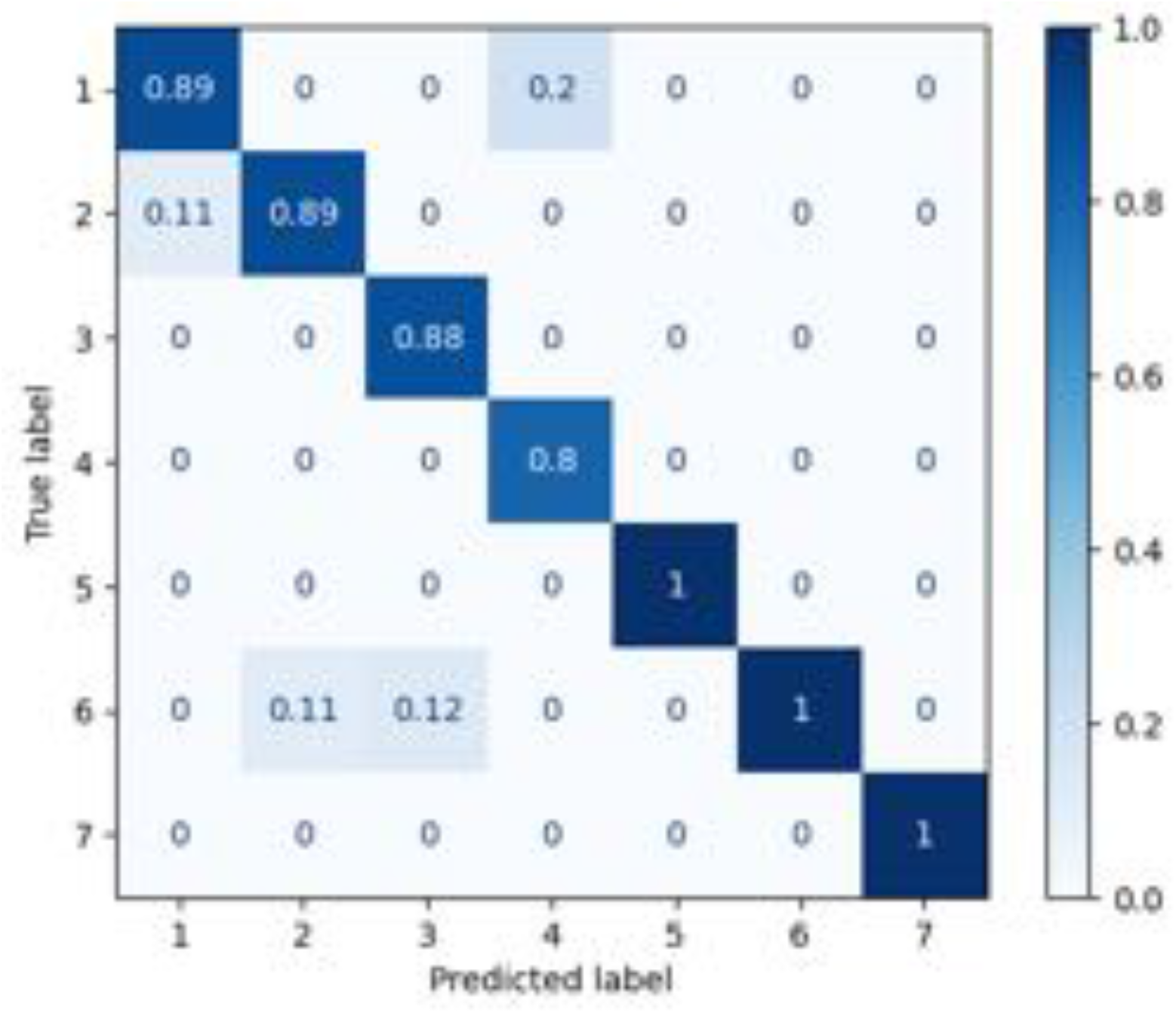

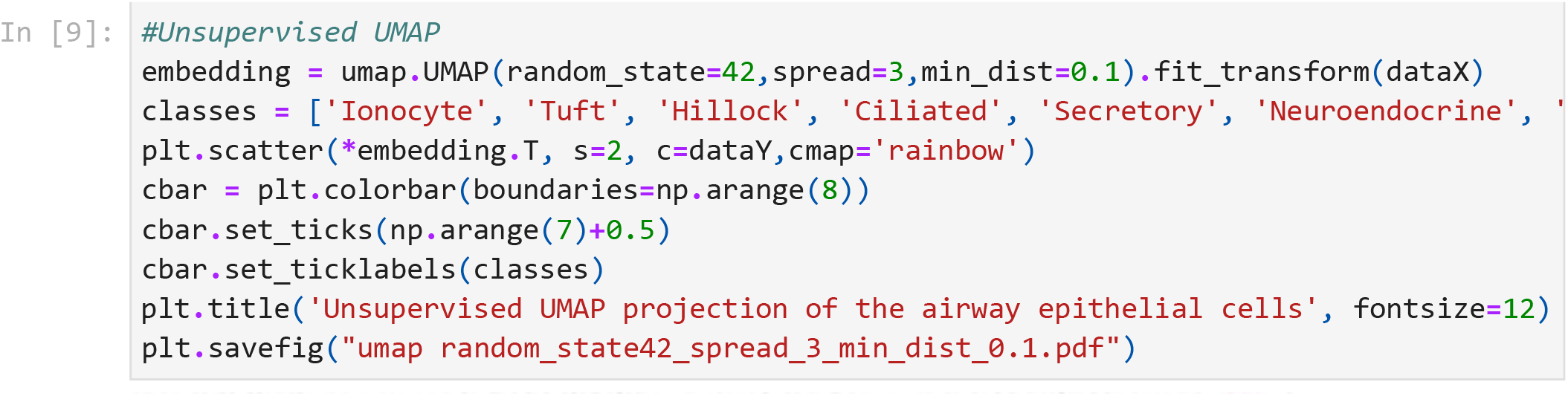

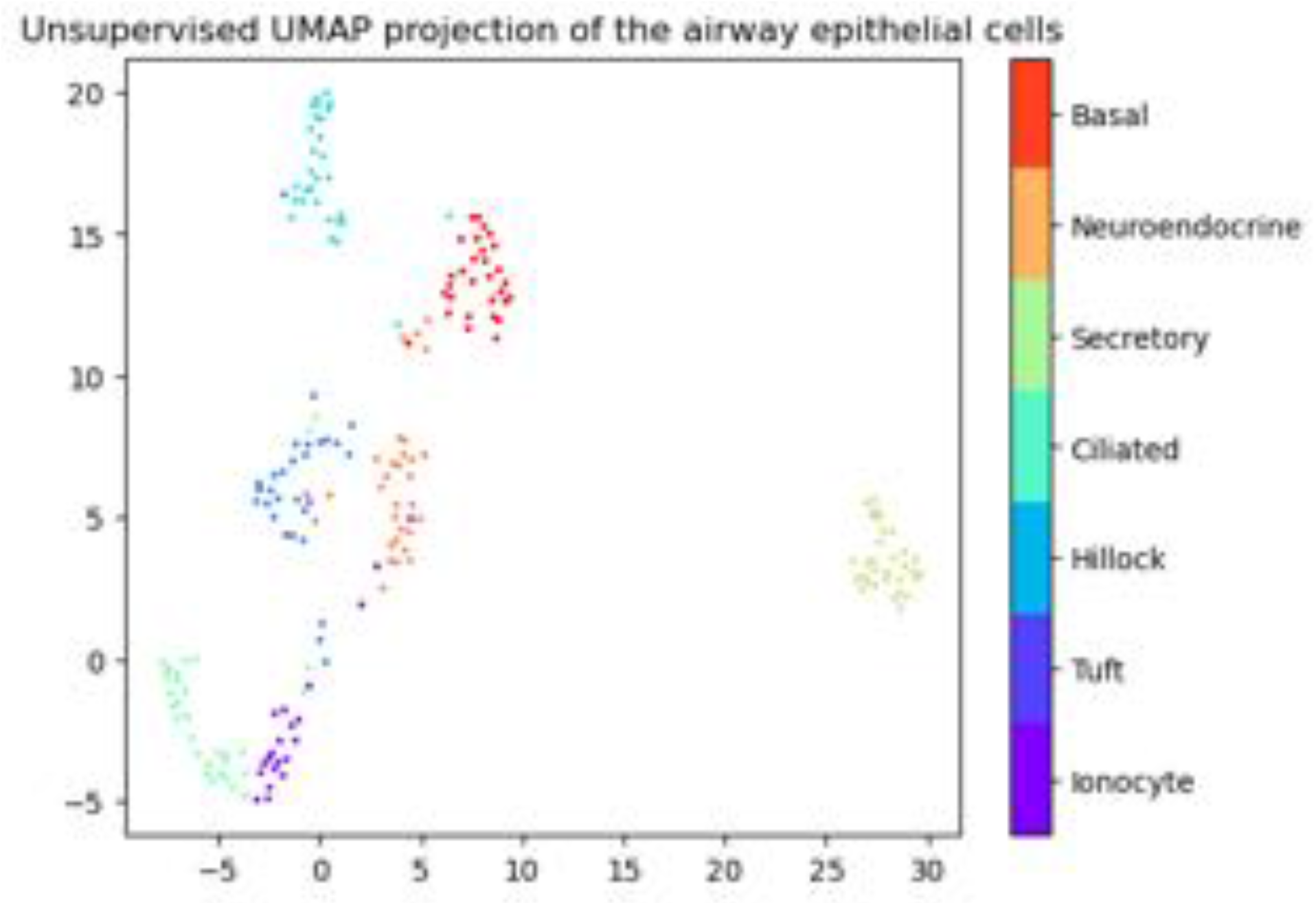

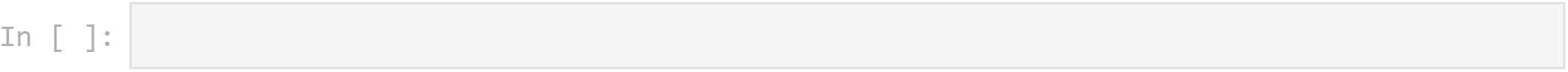

## References

Adler, K. B., Tuvim, M. J., & Dickey, B. F. (2013). Regulated Mucin Secretion from Airway Epithelial Cells. Frontiers in Endocrinology, 4(SEP). https://doi.org/10.3389/FENDO.2013.00129

Chance, B., Schoener, B., Oshino, R., Itshak, F., & Nakase, Y. (1979). Oxidation-reduction ratio studies of mitochondria in freeze-trapped samples. NADH and flavoprotein fluorescence signals. Journal of Biological Chemistry, 254(11), 4764–4771. https://doi.org/10.1016/S0021-9258(17)30079-0

Deprez, M., Zaragosi, L. E., Truchi, M., Becavin, C., García, S. R., Arguel, M. J., Plaisant, M., Magnone, V., Lebrigand, K., Abelanet, S., Brau, F., Paquet, A., Pe’er, D., Marquette, C. H., Leroy, S., & Barbry, P. (2020). A Single-Cell Atlas of the Human Healthy Airways. American Journal of Respiratory and Critical Care Medicine, 202(12), 1636–1645. https://doi.org/10.1164/RCCM.201911-2199OC

Dilipkumar, A., Al-Shemmary, A., Kreiß, L., Cvecek, K., Carlé, B., Knieling, F., Gonzales Menezes, J., Thoma, O. M., Schmidt, M., Neurath, M. F., Waldner, M., Friedrich, O., & Schürmann, S. (2019). Label-Free Multiphoton Endomicroscopy for Minimally Invasive In Vivo Imaging. Advanced Science (Weinheim, Baden-Wurttemberg, Germany), 6(8). https://doi.org/10.1002/ADVS.201801735

Fischer, A. J., Pino-Argumedo, M. I., Hilkin, B. M., Shanrock, C. R., Gansemer, N. D., Chaly, A. L., Zarei, K., Allen, P. D., Ostedgaard, L. S., Hoffman, E. A., Stoltz, D. A., Welsh, M. J., & Abou Alaiwa, M. H. (2019). Mucus strands from submucosal glands initiate mucociliary transport of large particles. JCI Insight, 4(1), 124863. https://doi.org/10.1172/JCI.INSIGHT.124863

Gil, D. A., Sharick, J. T., Mancha, S., Gamm, U. A., Choma, M. A., & Skala, M. C. (2019). Redox imaging and optical coherence tomography of the respiratory ciliated epithelium. Journal of Biomedical Optics, 24(1), 1. https://doi.org/10.1117/1.JBO.24.1.010501

Goldfarbmuren, K. C., Jackson, N. D., Sajuthi, S. P., Dyjack, N., Li, K. S., Rios, C. L., Plender, E. G., Montgomery, M. T., Everman, J. L., Bratcher, P. E., Vladar, E. K., & Seibold, M. A. (2020). Dissecting the cellular specificity of smoking effects and reconstructing lineages in the human airway epithelium. Nature Communications, 11(1). https://doi.org/10.1038/s41467-020-16239-z

Gustafsson, J. K., Davis, J. E., Rappai, T., McDonald, K. G., Kulkarni, D. H., Knoop, K. A., Hogan, S. P., Fitzpatrick, J. A. J., Lencer, W. I., & Newberry, R. D. (2021). Intestinal goblet cells sample and deliver lumenal antigens by regulated endocytic uptake and transcytosis. ELife, 10. https://doi.org/10.7554/ELIFE.67292

Gustafsson, J. K., & Johansson, M. E. V. (2022). The role of goblet cells and mucus in intestinal homeostasis. Nature Reviews Gastroenterology & Hepatology 2022, 1–19. https://doi.org/10.1038/s41575-022-00675-x

Heikal, A. A. (2010). Intracellular coenzymes as natural biomarkers for metabolic activities and mitochondrial anomalies. Biomarkers in Medicine, 4(2), 241. https://doi.org/10.2217/BMM.10.1

Hou, J., Wright, H. J., Chan, N., Tran, R., Razorenova, O. v., Potma, E. O., & Tromberg, B. J. (2016). Correlating two-photon excited fluorescence imaging of breast cancer cellular redox state with seahorse flux analysis of normalized cellular oxygen consumption. Journal of Biomedical Optics, 21(6), 060503. https://doi.org/10.1117/1.JBO.21.6.060503

Huang, S., Heikal, A. A., & Webb, W. W. (2002). Two-photon fluorescence spectroscopy and microscopy of NAD(P)H and flavoprotein. Biophysical Journal, 82(5), 2811–2825. https://doi.org/10.1016/S0006-3495(02)75621-X

Kasischke, K. A., Vishwasrao, H. D., Fisher, P. J., Zipfel, W. R., & Webb, W. W. (2004). Neural activity triggers neuronal oxidative metabolism followed by astrocytic glycolysis. Science (New York, N.Y.), 305(5680), 99–103. https://doi.org/10.1126/SCIENCE.1096485

Katz, A., Savage, H. E., Schantz, S. P., McCormick, S. A., & Alfano, R. R. (2002). Noninvasive native fluorescence imaging of head and neck tumors. Technology in Cancer Research & Treatment, 1(1), 9–15. https://doi.org/10.1177/153303460200100102

Klinger, A., Orzekowsky-Schroeder, R., von Smolinski, D., Blessenohl, M., Schueth, A., Koop, N., Huettmann, G., & Gebert, A. (2012). Complex morphology and functional dynamics of vital murine intestinal mucosa revealed by autofluorescence 2-photon microscopy. Histochemistry and Cell Biology, 137(3), 269–278. https://doi.org/10.1007/S00418-011-0905-0

Knoop, K. A., McDonald, K. G., McCrate, S., McDole, J. R., & Newberry, R. D. (2014). Microbial sensing by goblet cells controls immune surveillance of luminal antigens in the colon. Mucosal Immunology 2015 8:1, 8(1), 198–210. https://doi.org/10.1038/mi.2014.58

Kreiß, L., Thoma, O. M., Dilipkumar, A., Carlé, B., Longequeue, P., Kunert, T., Rath, T., Hildner, K., Neufert, C., Vieth, M., Neurath, M. F., Friedrich, O., Schürmann, S., & Waldner, M. J. (2020). Label-Free In Vivo Histopathology of Experimental Colitis via 3-Channel Multiphoton Endomicroscopy. Gastroenterology, 159(3), 832–834. https://doi.org/10.1053/J.GASTRO.2020.05.081

Lemire, S., Thoma, O. M., Kreiss, L., Völkl, S., Friedrich, O., Neurath, M. F., Schürmann, S., & Waldner, M. J. (2022). Natural NADH and FAD Autofluorescence as Label-Free Biomarkers for Discriminating Subtypes and Functional States of Immune Cells. International Journal of Molecular Sciences, 23(4). https://doi.org/10.3390/IJMS23042338/S1

Lin, B., Vinarsky, V., Shah, V., Chernoff, C., Waghray, A., Xu, J., Surve, M., Sun, J., Capen, D., Villoria, J., Brown, D., Hariri, L., & Rajagopal, J. (2022). Airway Hillocks are Injury-Resistant Reservoirs of Plastic Stem Cells. Science, Submitted.

Massaro, G. D., Paris, M., & Aung Thet, L. (1979). In vivo regulation of secretion of bronchiolar Clara cells in rats. The Journal of Clinical Investigation, 63(2), 167–172. https://doi.org/10.1172/JCI109285

McDole, J. R., Wheeler, L. W., McDonald, K. G., Wang, B., Konjufca, V., Knoop, K. A., Newberry, R. D., & Miller, M. J. (2012). Goblet cells deliver luminal antigen to CD103+ dendritic cells in the small intestine. Nature 2012 483:7389, 483(7389), 345–349. https://doi.org/10.1038/nature10863

Montoro, D. T., Haber, A. L., Biton, M., Vinarsky, V., Lin, B., Birket, S. E., Yuan, F., Chen, S., Leung, H. M., Villoria, J., Rogel, N., Burgin, G., Tsankov, A. M., Waghray, A., Slyper, M., Waldman, J., Nguyen, L., Dionne, D., Rozenblatt-Rosen, O., … Rajagopal, J. (2018). A revised airway epithelial hierarchy includes CFTR-expressing ionocytes. Nature, 560(7718), 319–324. https://doi.org/10.1038/s41586-018-0393-7

Mou, H., Vinarsky, V., Tata, P. R., Brazauskas, K., Choi, S. H., Crooke, A. K., Zhang, B., Solomon, G. M., Turner, B., Bihler, H., Harrington, J., Lapey, A., Channick, C., Keyes, C., Freund, A., Artandi, S., Mense, M., Rowe, S., Engelhardt, J. F., … Rajagopal, J. (2016). Dual SMAD Signaling Inhibition Enables Long-Term Expansion of Diverse Epithelial Basal Cells. Cell Stem Cell, 19(2), 217–231. https://doi.org/10.1016/J.STEM.2016.05.012

Noah, T. K., Knoop, K. A., McDonald, K. G., Gustafsson, J. K., Waggoner, L., Vanoni, S., Batie, M., Arora, K., Naren, A. P., Wang, Y. H., Lukacs, N. W., Munitz, A., Helmrath, M. A., Mahe, M. M., Newberry, R. D., & Hogan, S. P. (2019). IL-13-induced Intestinal secretory epithelial cell antigen passages are required for IgE-mediated food-induced anaphylaxis. The Journal of Allergy and Clinical Immunology, 144(4), 1058. https://doi.org/10.1016/J.JACI.2019.04.030

Pilette, C., Godding, V., Kiss, R., Delos, M., Verbeken, E., Decaestecker, C., de Paepe, K., Vaerman, J. P., Decramer, M., & Sibille, Y. (2012). Reduced Epithelial Expression of Secretory Component in Small Airways Correlates with Airflow Obstruction in Chronic Obstructive Pulmonary Disease. https://Doi.Org/10.1164/Ajrccm.163.1.9912137, 163(1), 185-194. https://doi.org/10.1164/AJRCCM.163.1.9912137

Plasschaert, L. W., Žilionis, R., Choo-Wing, R., Savova, V., Knehr, J., Roma, G., Klein, A. M., & Jaffe, A. B. (2018). A single-cell atlas of the airway epithelium reveals the CFTR-rich pulmonary ionocyte. In Nature (Vol. 560, Issue 7718, pp. 377–381). Nature Publishing Group. https://doi.org/10.1038/s41586-018-0394-6

Rocheleau, J. v., Head, W. S., & Piston, D. W. (2004). Quantitative NAD(P)H/flavoprotein autofluorescence imaging reveals metabolic mechanisms of pancreatic islet pyruvate response. The Journal of Biological Chemistry, 279(30), 31780–31787. https://doi.org/10.1074/JBC.M314005200

Schaefer, P. M., Kalinina, S., Rueck, A., von Arnim, C. A. F., & von Einem, B. (2019). NADH Autofluorescence-A Marker on its Way to Boost Bioenergetic Research. Cytometry Part A, 95(1), 34–46. https://doi.org/10.1002/CYTO.A.23597

Sepehr, R., Staniszewski, K., Maleki, S., Jacobs, E. R., Audi, S., & Ranji, M. (2012). Optical imaging of tissue mitochondrial redox state in intact rat lungs in two models of pulmonary oxidative stress. Journal of Biomedical Optics, 17(4), 046010. https://doi.org/10.1117/1.JBO.17.4.046010

Shijubo, N., Itoh, Y., Yamaguchi, T., Imada, A., Hirasawa, M., Yamada, T., Kawai, T., & Abe, S. (2012). Clara Cell Protein-positive Epithelial Cells Are Reduced in Small Airways of Asthmatics. https://Doi.Org/10.1164/Ajrccm.160.3.9803113, 160(3), 930-933. https://doi.org/10.1164/AJRCCM.160.3.9803113

Shijubo, N., Itoh, Y., Yamaguchi, T., Shibuya, Y., Morita, Y., Hirasawa, M., Okutani, R., Kawai, T., & Abe, S. (1997). Serum and BAL Clara cell 10 kDa protein (CC10) levels and CC10-positive bronchiolar cells are decreased in smokers. Eur Respir J, 10, 1108–1114. https://doi.org/10.1183/09031936.97.10051108

Skala, M. C., Riching, K. M., Gendron-Fitzpatrick, A., Eickhoff, J., Eliceiri, K. W., White, J. G., & Ramanujam, N. (2007). In vivo multiphoton microscopy of NADH and FAD redox states, fluorescence lifetimes, and cellular morphology in precancerous epithelia. Proceedings of the National Academy of Sciences of the United States of America, 104(49), 19494–19499. https://doi.org/10.1073/PNAS.0708425104/ASSET/83A234B6-C96C-4A70-AB00-07E97F6DE60D/ASSETS/GRAPHIC/ZPQ0480784940004.JPEG

Tiede, L. M., Rocha-Sanchez, S. M., Hallworth, R., Nichols, M. G., & Beisel, K. (2007). Determination of hair cell metabolic state in isolated cochlear preparations by two-photon microscopy. Journal of Biomedical Optics, 12(2), 021004. https://doi.org/10.1117/1.2714777

Webber, S. E., & Widdicombe, J. G. (1987). The actions of methacholine, phenylephrine, salbutamol and histamine on mucus secretion from the ferret in-vitro trachea. Agents and Actions, 22(1-2), 82–85. https://doi.org/10.1007/BF01968821

Wong, A. P., Keating, A., & Waddell, T. K. (2009). Airway regeneration: the role of the Clara cell secretory protein and the cells that express it. Cytotherapy, 11(6), 676–687. https://doi.org/10.3109/14653240903313974

Yoneda, K. (1977). Pilocarpine stimulation of the bronchiolar Clara cell secretion. Laboratory Investigation; a Journal of Technical Methods and Pathology, 37(5), 447–452. https://europepmc.org/article/med/916620

